# Molecular basis of AvrRvi6 recognition and escape from *Rvi6*-mediated immunity

**DOI:** 10.64898/2026.11.26.769449

**Authors:** Sirine Benmamar, Mélanie Sannier, Bruno Le Cam, Maël Baudin

## Abstract

Apple scab, caused by the fungal pathogen *Venturia inaequalis*, is the major disease of apple orchards. The receptor-like protein *Rvi6* confers resistance against *V inaequalis* isolates carrying the cognate avirulence effector *AvrRvi6*. AvrRvi6 is the first apoplastic MAX effector described, providing a unique system to study extracellular recognition of this fungal effector family. Here, using the heterologous system *Nicotiana benthamiana,* we investigated the molecular basis of AvrRvi6 recognition by Rvi6 using structural modeling, protein–protein interaction assays, amino acids substitutions, and live-cell imaging. We show that *Rvi6*-mediated hypersensitive response requires the receptor-like kinase co-receptors *SOBIR1* and *BAK1*. Natural AvrRvi6 variants study revealed that virulent variants displayed an altered subcellular localization. In addition, protein complex structural modeling identified a putative extracellular interface between AvrRvi6 and Rvi6 involving multiple contact surfaces. Yeast two-hybrid assays supported their association while reciprocal mutagenesis of predicted interface residues impaired or abolished Rvi6-mediated recognition, providing functional support for their contribution to effector recognition. Together, our results support a model in which Rvi6 recognizes AvrRvi6 through a multivalent extracellular interface and suggest that some virulent alleles evade immune surveillance through altered protein trafficking. These findings provide mechanistic insight into the extracellular effector recognition and reveal how naturally occurring variation in a fungal MAX effector can modulate host immune recognition.

## Introduction

Apple scab, caused by the Ascomycota fungus *Venturia inaequalis*, is the most economically devastating disease affecting apple orchards worldwide (Bowen et al. 2011). Upon infection, spores germinate and penetrate the plant cuticle, where the fungus adopts a highly specialized lifestyle by developing pseudoparenchymatous stromata and runner hyphae entirely within the subcuticular space. The pathogen does not penetrate the underlying epidermal cells, maintaining a restricted extracellular niche (Steiner and Oerke 2024). During the colonization of apple tissues, *V. inaequalis* expresses successive waves of hundreds of small secreted proteins, called effectors (Rocafort et al. 2022). These proteins can be classified into several structural families and are hypothesized to play a central role in fungal virulence, effectively suppressing basal defenses to promote virulence (Deng et al. 2017; Rocafort et al. 2022, 2023).

To detect these extracellular threats, plants have evolved a sophisticated immune surveillance system governed by plasma membrane-localized Pattern Recognition Receptors (PRRs) that monitor the apoplast for invasion patterns and effectors. A major class of PRRs involved in this defense are Receptor-Like Proteins (RLPs) (Stotz et al. 2014). RLPs are characterized by an extracellular Leucine-Rich Repeat (LRR) domain responsible for ligand binding, a transmembrane domain, and a short cytoplasmic tail that notably lacks an intracellular kinase domain (Albert et al. 2015). These receptors have been found to be involved in gene-for-gene resistance against many apoplastic fungal pathogens such as the tomato Cf proteins Cf-4 and Cf-9 that recognize the *Cladosporium fulvum* effectors Avr4 and Avr9 respectively (Kruijt et al. 2005; Thomas et al. 1995; Wulff et al. 2001). Because RLPs lack their own intracellular signaling capacity, they require the recruitment of Receptor-Like Kinase (RLK) co-receptors to transmit immune signals into the cell. RLPs usually associate with the regulatory RLK SOBIR1 (Suppressor of BIR1-1) in a ligand-independent manner (Gust and Felix 2014; Liebrand et al. 2013). Upon direct or indirect perception of their cognate apoplastic effector, this complex recruits an additional co-receptor, most notably the SOMATIC EMBRYOGENESIS RECEPTOR KINASE (SERK) BAK1 (BRI1-Associated Kinase 1) (Liebrand et al. 2014; Albert et al. 2015). The formation of this highly active complex triggers transphosphorylation events that initiate intracellular signaling cascades, ultimately culminating in a form of programmed cell death called hypersensitive response (HR) and the restriction of pathogen growth. An example of this mechanism is the recognition of Avr4 by Cf-4 aforementioned, in which BAK1 (Postma et al. 2016) and SOBIR1 (Liebrand et al. 2013) are required for the induction of the immune response.

In the *Malus–Venturia* pathosystem, the *Rvi6* gene (formerly *HcrVf2*) encodes an RLP that confers strong resistance to apple scab (Belfanti et al. 2004). Recently, *AvrRvi6,* an apoplastic effector secreted by *V. inaequalis* was identified as the avirulence determinant triggering *Rvi6* immunity (Sannier et al. 2025). Genetic variation conferring virulence at this effector locus arose thousands of years before apple domestication (Sannier et al. 2025). AvrRvi6 remarkably adopts a MAX (*Magnaporthe* Avrs and ToxB-like) structural fold, a beta-sandwich topology originally characterized in cytoplasmic effectors of the rice blast fungus *Magnaporthe oryzae* (De Guillen et al. 2015; Naour-Vernet et al. 2025). The MAX effector family is the largest effector family identified in *V. inaequalis*, comprising 75 predicted members (Rocafort et al. 2022). The recognition of AvrRvi6 by Rvi6 at the cell surface triggers an allele-specific HR (Sannier et al. 2025). However, natural populations of *V. inaequalis* have managed to circumvent this resistance through various evolutionary mechanisms, including partial sequence deletions, transposon insertions in the promoter leading to a loss of expression, and non-synonymous mutations in the coding sequence (Sannier et al. 2025). Despite the identification of Rvi6 and AvrRvi6, the precise structural dynamics governing their interaction and the assembly of the immune complex remain to be fully elucidated. In this study, we deciphered the structural basis underlying the recognition of the *V. inaequalis* effector AvrRvi6 by the apple immune receptor Rvi6. Using structural modeling in combination with the heterologous systems *Nicotiana benthamiana* and yeast, we predicted and functionally validated residues involved in the association between AvrRvi6 and Rvi6. Using laser-scanning confocal microscopy, we investigated the subcellular co-localization of Rvi6 and AvrRvi6 and provided evidence that some virulent AvrRvi6 alleles may evade *Rvi6*-mediated immunity through improper secretion and altered subcellular localization. Together, our results provide structural and cellular insights into Rvi6–AvrRvi6 recognition and reveal a potential mechanism by which *V. inaequalis* escapes host immune surveillance.

## Results

### 1. BAK1 and SOBIR1 are required for AvrRvi6 recognition by Rvi6

To better understand the immune responses triggered by AvrRvi6, we first sought to characterize the downstream signaling pathway activated by its cognate receptor Rvi6. Leucine-rich repeat receptor-like proteins (LRR-RLPs), such as Rvi6 (Švara et al. 2024; Vinatzer et al. 2001), typically recruit the co-receptors BAK1 and SOBIR1 to initiate immune signaling (Gust and Felix 2014; Liebrand et al. 2014). To determine whether these co-receptors are required for Rvi6-mediated immunity, we utilized the *N. benthamiana* model system, in which we previously demonstrated that Rvi6 recognition of AvrRvi6 triggers a hypersensitive response (HR; Sannier et al. 2025). We tested Rvi6 signaling in *N. benthamiana bak1* and *sobir1* mutant backgrounds (Huang et al. 2021; Zönnchen et al. 2022). Transient co-expression of Rvi6 and AvrRvi6 from the EU-B04 strain (hereafter: AvrRvi6^EU-B04^) in the *bak1* mutant resulted in a complete loss of HR, whereas robust HR induction was observed in wild-type (WT) plants (Figure 1A). Quantification of HR phenotypes by red-light imaging (Figure 1B; Landeo Villanueva et al. 2021) confirmed that HR intensity was significantly attenuated in the *bak1* background compared to WT, demonstrating the requirement of BAK1 for AvrRvi6-triggered immunity in *N. benthamiana*. Similarly, co-expression of Rvi6 and AvrRvi6^EU-B04^ in two independent *sobir1* knockout lines (*sobir1-1* and *sobir1-2*) abolished the HR phenotype while eliciting normal HR in WT plants (Figure 1C). As a control, we monitored the Cf-4 and Avr4 pair, whose recognition depends on both BAK1 and SOBIR1 (Liebrand et al. 2013; Postma et al. 2016). The co-expression of Cf-4 and Avr4 induced HR in WT, the *bak1* line and *MdSOBIR1* complemented *sobir1* mutants, it failed to do so in uncomplemented *sobir1* mutants (Figure 1). Furthermore, transgenic expression of *Malus × domestica* SOBIR1 (MdSOBIR1) in the *sobir1* mutants fully restored the HR phenotype for both Rvi6 - AvrRvi6^EU-B04^ and Cf-4 - Avr4, as validated by HR quantification (Figures 1C, 1D). However, MdSOBIR1 expression induced a reduction of HR signals upon expression of Rvi6 and AvrRvi6^EU-B04^ in WT (Figures 1C, 1D). Laser-scanning confocal microscopy confirmed the plasma membrane localization of MdSOBIR1 (Figure S1), consistent with the localization reported for tomato SlSOBIR1 (Liebrand et al. 2013), although without a pronounced vesicular pattern. Taken together, these results demonstrate that *Rvi6*-mediated HR strictly requires BAK1 and SOBIR1 in *N. benthamiana*, and that apple MdSOBIR1 functionally complements *Nbsobir1* mutants.

**Figure 1.**
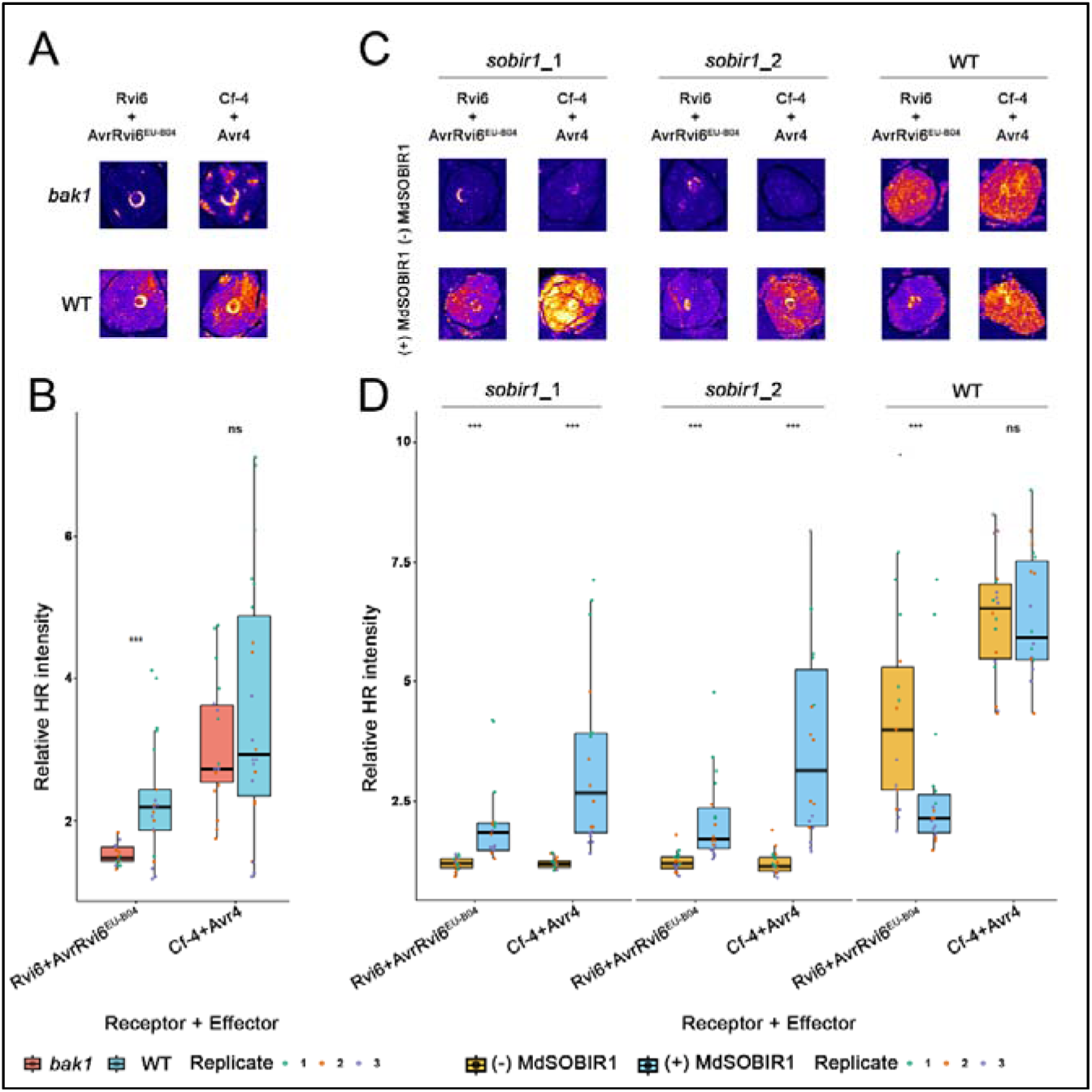
BAK1 and SOBIR1 are required for Rvi6-mediated recognition of AvrRvi6 in Nicotiana benthamiana. Hypersensitive response (HR) intensity (relative median value per infiltration spot) was measured using red-light imaging (Landeo Villanueva et al. 2021) for Cf-4 - Avr4 (control) and Rvi6 - AvrRvi6^EU-B04^ combinations transiently expressed via agroinfiltration. A) Phenotypic responses upon transient expression of the combinations in bak1 mutant or wild-type (WT). B) Quantification of the HR responses in (A) : bak1 (red) and wild-type (blue) genotypes; significance was determined using a two-tailed Student’s t-test or a Wilcoxon rank-sum test for non-normal data. C) Phenotypic responses upon transient expression of the combinations in two sobir1 mutants and WT with and without an MdSOBIR1 complementation. D) Quantification of HR responses in (C) faceted by Genotype. With (blue) and without (yellow) MdSOBIR1 complementation. Significance was determined via a Linear Mixed-Effects Model (LMM; fixed effects: Genotype × Unit × MdSOBIR1; random effects: ID_Plante and Replicate) followed by post-hoc pairwise emmeans comparisons. Points represent individual spots color-coded by replicate 3 independent biological replicates comprising 6 individual measures each were performed. Asterisks denote significant differences: * p < 0.05, p < 0.01, *** p < 0.001, ns (not significant). All HR intensity signals are relative to non-infiltrated-leaf signal intensity.

### 2. AvrRvi6 natural polymorphisms do not critically impact protein folding

We previously identified 20 natural allelic variants of AvrRvi6 in *V. inaequalis* populations that exhibit distinct capacities to trigger *Rvi6*-mediated immunity in *N. benthamiana* and correlate with virulence or avirulence on *Rvi6*-carrying apple cultivars (Sannier et al. 2025). To evaluate whether natural polymorphisms compromise recognition by destabilizing the effector’s tertiary structure, we predicted the structures of these variants using AlphaFold2 (Jumper et al. 2021). Structural alignments of AvrRvi6 alleles relative to the reference AvrRvi6^EU-B04^ revealed that the global protein architecture is conserved, with all pairwise root-mean-square deviation (RMSD) values remaining low (<0.5 Å; the truncated variant AvrRvi6^2828^ was excluded from global alignment) (Figure 2A), Despite this global conservation, minor local secondary structure variations were observed: All the virulent alleles and the avirulent allele AvrRvi6^1402^ harbor a slightly shorter α3 helix (in which C92 is excluded). In addition to this structural divergence, the virulent allele AvrRvi6^1153^ lacks the short α2 helix (residues 75–77). Structural dendrogram clustering and RMSD-based secondary structure comparisons identified the virulent AvrRvi6^1153^ and AvrRvi6^1180^ alleles as basal outliers diverging furthest from AvrRvi6^EU-B04^ (Figures 2A and S3). Mapping natural single-nucleotide polymorphisms (SNPs) onto the predicted 3D structure revealed that virulence-associated residues do not localize to the misaligned regions (Figure 2B). Similarly, shared polymorphic sites between virulent and avirulent alleles occur primarily in invariant structural regions; the only exception is T76S, which lies within the α2 helix present in all alleles except AvrRvi6^1153^. Calculating the solvent-accessible surface area (SASA) indicated that virulence-associated mutations target distinct structural environments (Figure 2C): C21S (AvrRvi6^2917^) and C50R (AvrRvi6^1153^) polymorphisms are located in fully buried core residues, whereas F36S (AvrRvi6^1180^) affects a slightly exposed residue. Altogether, these structural analyses demonstrate that natural polymorphisms associated with virulence allow *V. inaequalis* to evade *Rvi6* surveillance without causing major structural rearrangements or disrupting of the overall AvrRvi6 fold.

**Figure 2:**
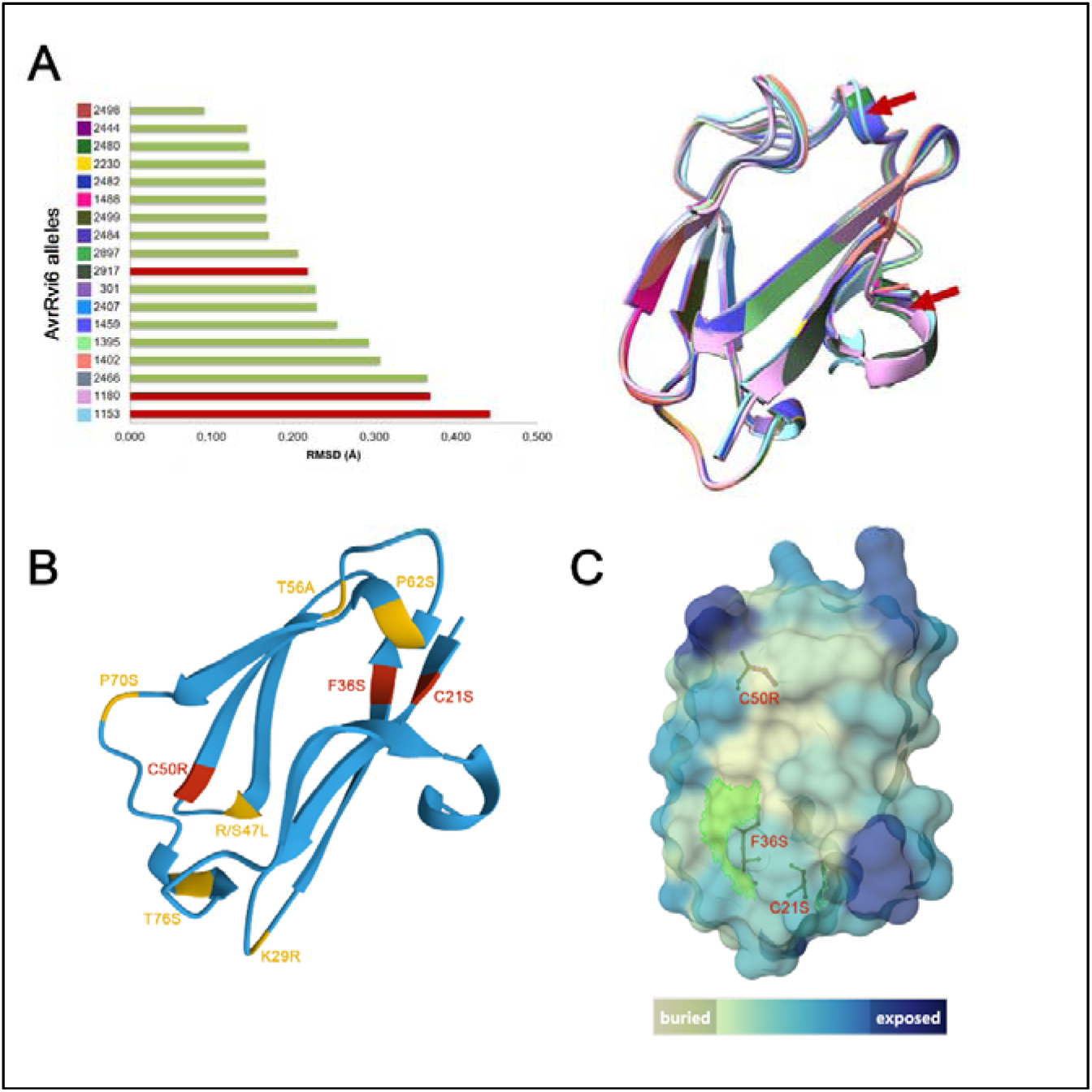
AvrRvi6 natural polymorphism does not critically impact protein folding. A) AvrRvi6 alleles’ structural alignment through ChimeraX with RMSD analysis. AvrRvi6^EU-B04^ is used as a reference. Red bars represent virulent alleles, green bars represent avirulent ones. Red arrows on the alignment point to structural divergences. B) Mapping of AvrRvi6 natural polymorphism on AlphaFold2 structural prediction. Red = SNPs present in virulent alleles only. Yellow = SNPs present in both virulent and avirulent alleles. C) AvrRvi6 3D protein modeling colored by solvent-accessible surface area (SASA): from beige for buried (low SASA) to dark blue for exposed (high SASA). SNPs linked to virulence are modelled in red ball and stick and identified, their presence/absence on the surface is shown by the presence/absence of green highlighting.

### 3. Virulent AvrRvi6 alleles do not accumulate in *N. benthamiana* apoplast

Since the virulent AvrRvi6 polymorphisms do not critically impact the protein structure, we then wondered if the mutations could affect the protein localization. To determine the subcellular localization of virulent AvrRvi6 alleles, we transiently co-expressed two virulent alleles (AvrRvi6^1180^ and AvrRvi6^1153^) alongside the reference avirulent allele AvrRvi6^EU-B04^ in *N. benthamiana* leaves. Constructs were expressed under the control of a double 35S promoter and co-infiltrated with the plasma membrane marker PITG16242 (an RXLR effector; Petre et al. 2021). To ensure targeting to the plant apoplast, the native fungal secretion signal of each variant was replaced with the plant PR1α signal peptide (PR1α^SP^), and an mCherry fluorescent reporter was fused to the C-terminus (PR1α^SP^-AvrRvi6^ΔSP^-mCherry). Laser-scanning confocal microscopy of *N. benthamiana* leaf epidermal cells at 3 days post-infiltration (dpi) showed that the avirulent fusion PR1α^SP^-AvrRvi6^EU-B04ΔSP^-mCherry localized predominantly to the apoplast, consistent with previous observations (Figure 3A; Sannier et al. 2025). In contrast, both virulent variants PR1α^SP^-AvrRvi6^1180ΔSP^-mCherry and PR1α^SP^-AvrRvi6^1153ΔSP^-mCherry accumulated intracellularly, displaying distinct fluorescence signals in vacuoles (Figures 3B and 3C). To validate these localization differences biochemically, we analyzed apoplastic wash fluids (AF) from the agroinfiltrated leaves by western blotting. The avirulent AvrRvi6^EU-B04ΔSP^-mCherry protein was readily detected in the apoplastic fraction (Figure 3D). In contrast, neither virulent variant was detectable in the AFs despite sharing the same plant-derived secretion signal (Figure 3D). Taken together, these results demonstrate that natural polymorphisms in virulent AvrRvi6 alleles impair effector accumulation in the apoplast and lead to intracellular retention, thereby preventing perception by the cell-surface receptor Rvi6.

**Figure 3:**
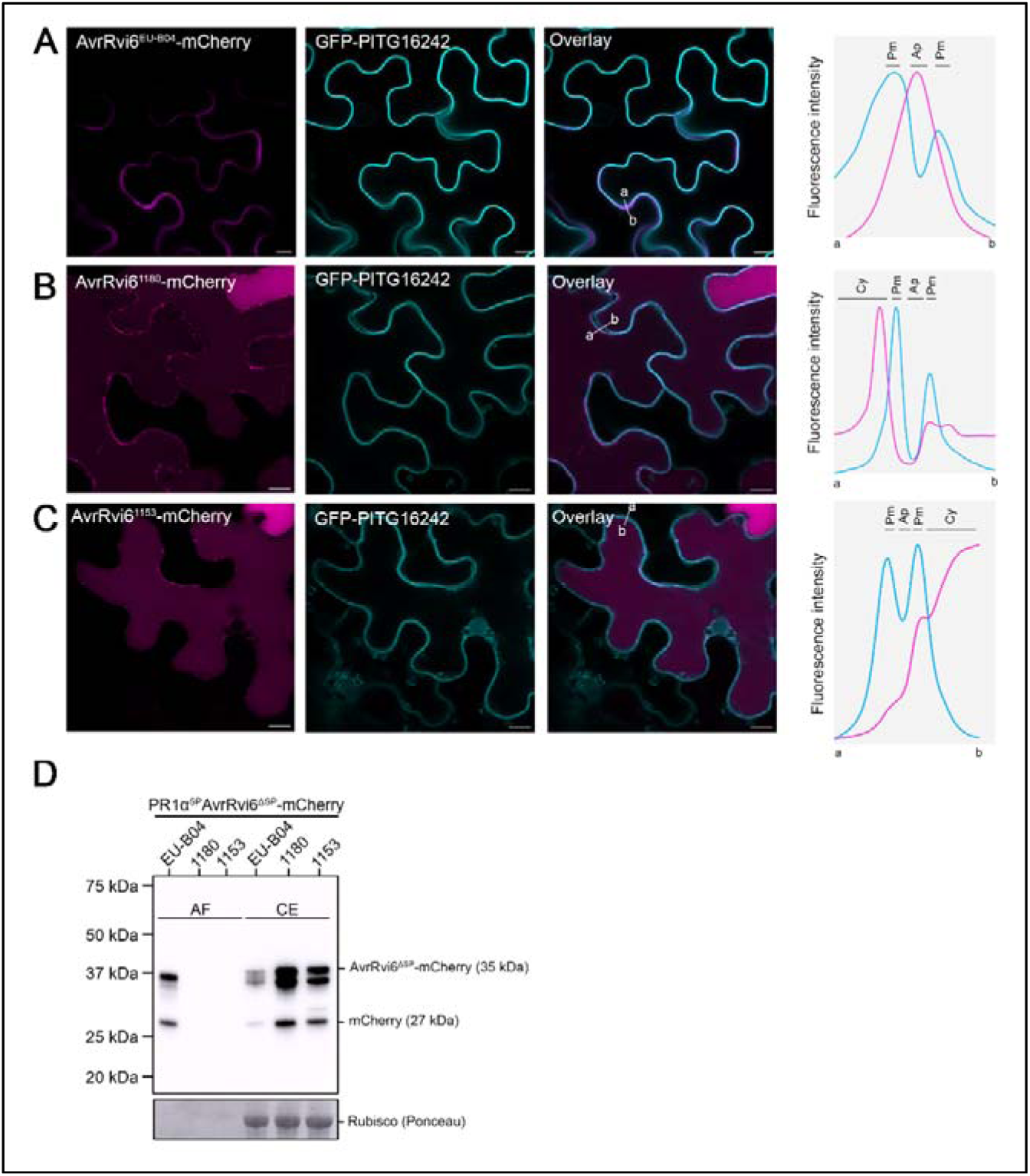
Virulent AvrRvi6 alleles are accumulated in the vacuole in *N. benthamiana*. (A–C) Subcellular localization of AvrRvi6 alleles in *N. benthamiana* leaf epidermal cells at 3 days post-infiltration (dpi) analyzed by laser-scanning confocal microscopy. Effector variants carrying the plant PR1α signal peptide and a C-terminal mCherry tag (PR1αSP-AvrRvi6ΔSP-mCherry magenta/red) were co-expressed under a double 35S promoter with the plasma membrane marker PITG16242-GFP (cyan; Petre et al. 2021). A) The avirulent variant EU-B04 localizes predominantly to the extracellular space (apoplast). (B, C) The virulent variants 1180 (B) and 1153 (C) fail to accumulate in the apoplast and instead localize intracellularly with distinct fluorescence signals in vacuoles. Pictures showing representative results from experiments that have been independently repeated at least twice. Scale bars =10 µm pm= plasma membrane, ap = apoplast, cy = cytosol. D) Immunoblot analysis of apoplastic wash fluids (AF) and total protein extracts (crude; CE) from *N. benthamiana* leaves transiently expressing the indicated constructs at 3 dpi. Effector proteins were detected using an anti-mCherry antibody. Apoplastic washing efficiency and cellular integrity were verified using ponceau staining.

### 4. Apoplastic AvrRvi6 relocalizes at the plasma membrane when co-expressed with Rvi6 in *N. benthamiana*

We previously established that the avirulent variant AvrRvi6^EU-B04^ accumulates in the apoplast (Figures 3A ; Sannier et al. 2025), a localization consistent with that of the immune receptor Rvi6 which is predominantly associated with the plasma membrane (Chen et al. 2025). To further validate the subcellular distribution of Rvi6, we transiently expressed Rvi6 C-terminally tagged with two distinct fluorescent reporters, mVenus and mStayGold, in *N. benthamiana*. Confocal imaging upon co-expression with the plasma membrane marker PITG16242-GFP (Petre et al. 2021) confirmed that Rvi6 localizes predominantly to the plasma membrane, with a fraction detectable in the endoplasmic reticulum network (Figures 4A and 4B). We next assessed whether the presence of Rvi6 alters the sub-cellular localization of AvrRvi6 *in planta*. In the absence of Rvi6 (expressed alongside the control marker PITG16242-GFP), AvrRvi6^EU-B04^ displayed its typical diffuse apoplastic distribution (Figure 4C). However, upon co-expression with Rvi6-mVenus, the AvrRvi6 fluorescence signal shifted considerably, concentrating at the cell periphery in close alignment with the plasma membrane (Figure 4D). Together, these observations demonstrate that the apoplastic AvrRvi6 is recruited to the plasma membrane in the presence of Rvi6, consistent with physical receptor–effector association at the host cell surface.

**Figure 4.**
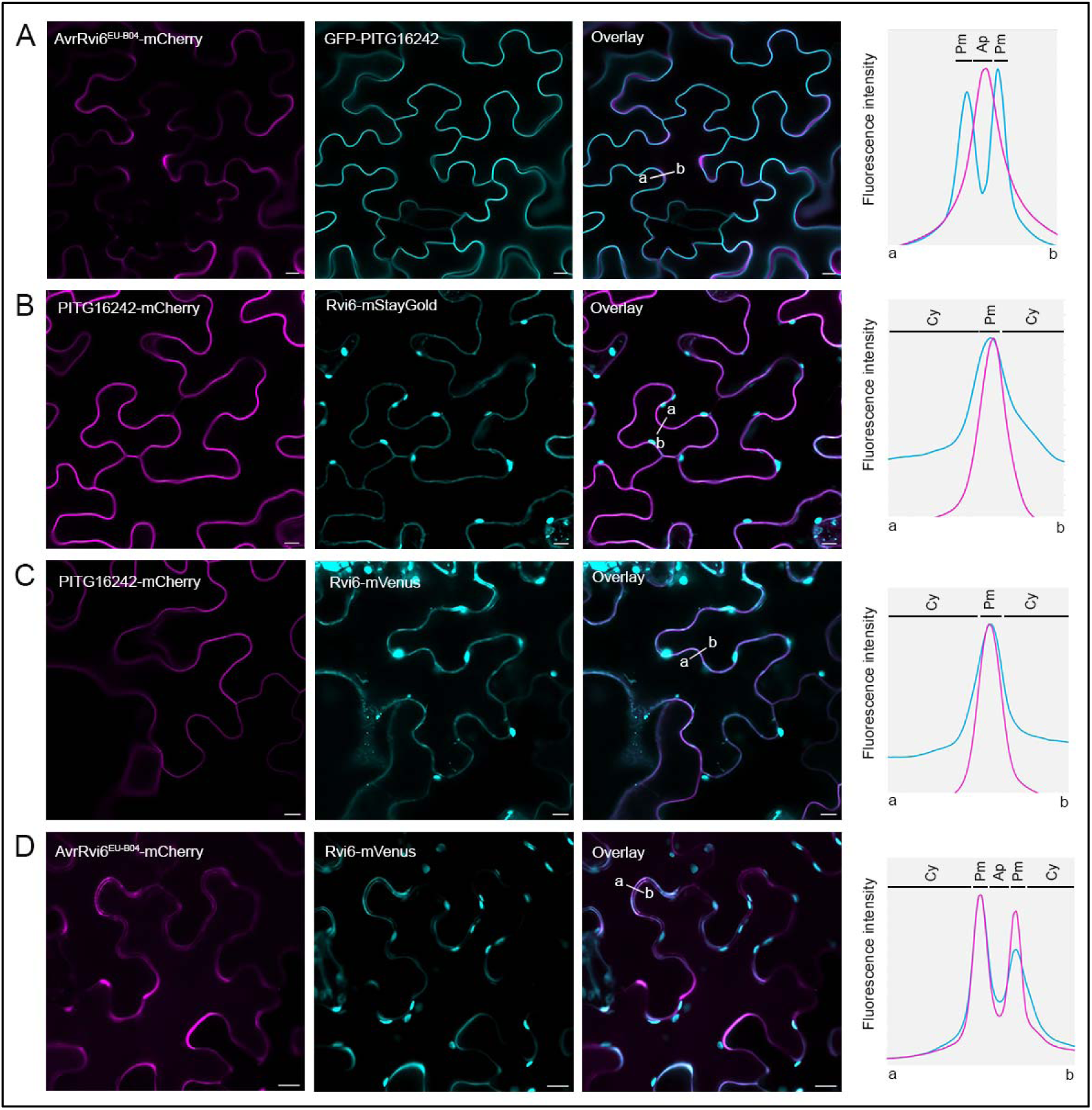
Apoplastic avirulent AvrRvi6 relocalizes to the plasma membrane in the presence of Rvi6 in *N. benthamiana*. (A-D) Subcellular localization of proteins in *N. benthamiana* leaf epidermal cells at 3 dpi analyzed by laser-scanning confocal microscopy. (A, B) Subcellular localization of Rvi6 tagged with mStayGold (A) or mVenus (B) (cyan) co-expressed with the plasma membrane marker PITG16242-mCherry (magenta; Petre et al. 2021). C) Subcellular localization of PR1α^SP^-AvrRvi6^EU-B04ΔSP^-mCherry (magenta) co-expressed with the plasma membrane marker PITG16242-GFP (cyan ; Petre et al. 2021). D) Subcellular localization of PR1α^SP^-AvrRvi6^EU-B04ΔSP^-mCherry (magenta) upon co-expression with Rvi6-mVenus (cyan). Pictures showing representative results from experiments that have been independently repeated at least twice. Scale bars = 10µm.

### 5. Structural modeling of Rvi6 in complex with AvrRvi6 identifies key residues for association

Given the apoplastic accumulation of AvrRvi6 (Sannier et al. 2025) and its recruitment to the plasma membrane upon co-expression with Rvi6 (Figure 4), we investigated the structural basis of direct receptor–effector association using AlphaFold-Multimer (Evans et al. 2021; Jumper et al. 2021; Figure 5). The predicted Rvi6 – AvrRvi6^EU-B04^ heterodimer yielded an overall model confidence score (0.8ipTM + 0.2pTM) of 0.53, which is below the standard high-confidence threshold of 0.70. Despite this moderate global metric, analysis of the predicted interface revealed a coherent, chemically complementary binding mode between both partners (Figure 5B). Thirteen AvrRvi6 residues were predicted to be involved in the association with Rvi6 (Figure 5B). Eleven Rvi6 residues were predicted to be participating in the association with AvrRvi6, with multiple ones involved in a cluster (Y94-F102) (Figure 5B). The predicted binding interface on Rvi6 is mainly located within solvent-exposed loop regions and is heavily enriched in aromatic residues, including phenylalanine (F102, F174 and F761), tryptophan (W97 and W651), and tyrosine (Y94, Y149 and Y175) (Figure 5B). These aromatic residues may contribute to hydrophobic interactions at the predicted interface. The interacting residues originate from two discontinuous sequence regions of Rvi6—an N-terminal-proximal cluster and a C-terminal region—that converge upon tertiary folding to form a claw-like interface that engages AvrRvi6 from opposing sides (Figure 5B). Complementing this electron-rich aromatic surface, the AvrRvi6 contact surface predominantly engages positively charged arginine residues (R65, R67, and R89). (Figure 5B). To determine whether predicted structural stability correlates with biological recognition, we generated complex models for all 20 natural AvrRvi6 allelic variants paired with Rvi6. Although complexes involving virulent alleles generally exhibited lower interface confidence scores than a subset of avirulent variants, their scores remained comparable to or higher than those of other avirulent alleles (Figure S4). Consequently, model confidence metrics alone do not quantitatively correlate with virulence across natural populations, suggesting that subtle local interface perturbations or intracellular trafficking defects govern immune evasion.

**Figure 5.**
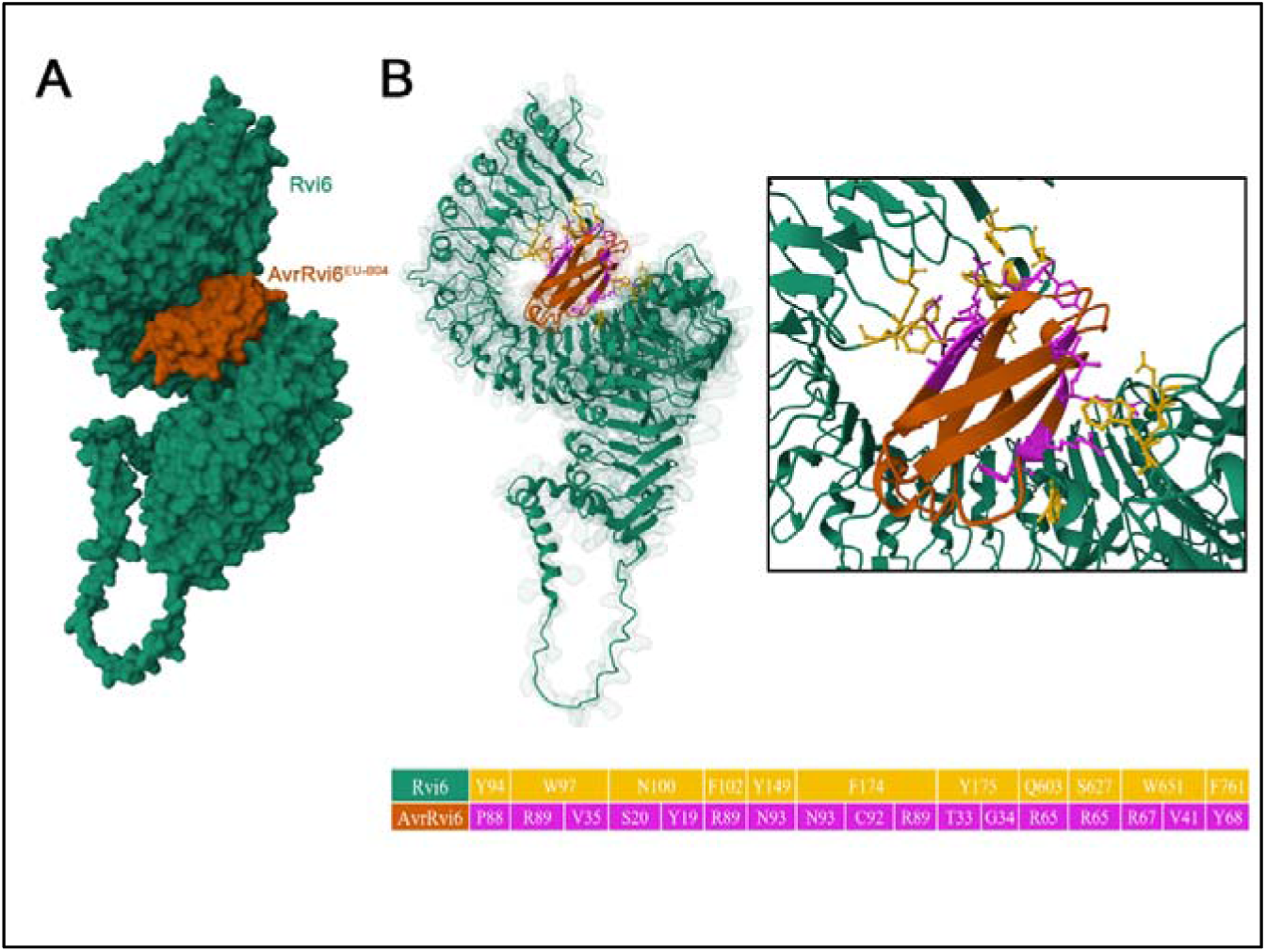
Structural modeling of Rvi6 in complex with AvrRvi6 identifies key residues for association. (A) Gaussian surface representation of the Rvi6–AvrRvi6^EU-B04^ heterodimeric complex predicted by AlphaFold-Multimer. Model confidence score (pLDDT=82.9 pTM=0.765 ipTM=0.472). (B) Secondary structures cartoon representation and detailed view of the predicted receptor–effector interface. Residues involved in the Rvi6 – AvrRvi6 recognition are shown in the table with Rvi6 residues on the top row linked to the AvrRvi6 residues in the bottom row. Color coding: Rvi6 in green with recognition sites residues in yellow and AvrRvi6 in dark orange with recognition site residues in pink.

### 5. AvrRvi6 associates with Rvi6 in yeast

We established that AvrRvi6 colocalizes with Rvi6 at the cell periphery (Figure 4) and that structural modeling predicts a direct receptor–effector binding interface (Figure 5). To experimentally test whether AvrRvi6 directly associates with Rvi6, we performed yeast two-hybrid (Y2H) assays. The mature forms (lacking native signal peptides, ΔSP) of both the avirulent AvrRvi6^EU-B04^ and virulent AvrRvi6^1180^ variants were fused to the GAL4 activation domain (GAL4-AD). These preys were tested against three bait truncations of Rvi6 fused to the GAL4 DNA-binding domain (GAL4-BD): Rvi6_LRR29–930_ (encompassing the complete extracellular LRR domain without the signal peptide), Rvi6_LRR45–633_ (harboring the core predicted AvrRvi6-binding interface), and Rvi6_LRR45–185_ (containing only some of the contact residues) (Figure 6). All co-transformed yeast strains grew on non-selective medium (SD-L-W; Figure 6A), and steady-state protein expression was confirmed by immunoblot analysis for all but T antigen (Figure S5A). On selective medium lacking histidine (SD-L-W-H), Rvi6_LRR45–633_ showed clear association with both AvrRvi6^EU-B04^ and the virulent variant AvrRvi6^1180^ (Figure 6B). Because structural alignments revealed that AvrRvi6 shares major structural homology with the *Magnaporthe oryzae* MAX (**M**agnaporthe **A**vr and To**x**B-like) effector family members AvrPikD and AvrPia, but less with Pwl2 (Maqbool et al. 2015; Ose et al. 2015; Sweigard et al. 1995; Figure S5B), we also tested whether Rvi6 interacts with unrelated MAX effectors. Rvi6_LRR45–633_ associated with AvrPikD^ΔSP^ and AvrPia^ΔSP^ on SD-L-W-H, but failed to associate with Pwl2^ΔSP^ (Figure 6B). In contrast, neither AvrRvi6 variant associated with the known NLR RGA5_HMA_ (Cesari et al. 2013; Ortiz et al. 2017) or with the negative control p53 (Li and Fields 1993), demonstrating interface specificity (Figure 6B). Under more stringent selection (SD-L-W-H supplemented with 0.1 mM 3-amino-1,2,4-triazole (3-AT)), robust yeast growth was exclusively maintained for the Rvi6_LRR45–633_ - AvrRvi6^EU-B04^ combination and positive controls (Figure 6C). In contrast, yeast expressing Rvi6_LRR45–633_ paired with AvrRvi6^1180^, AvrPia, or AvrPikD exhibited only weak residual growth under these conditions (Figure 6C). Given the association of *M. oryzae* MAX effectors with Rvi6 in yeast, we evaluated whether these heterologous effectors could trigger Rvi6-mediated defense in plant tissue. Transient co-expression of Rvi6 with PR1α^SP^-targeted AvrPikD, AvrPia, or Pwl2 in *N. benthamiana* failed to induce an HR, despite verified apoplastic effector accumulation for all but Pwl2 (Figures S5C and S5D). Collectively, these results demonstrate that Rvi6_LRR45–633_ selectively engages MAX-fold effectors in yeast, displaying a considerably higher association affinity for the avirulent AvrRvi6^EU-B04^ allele compared to its virulent counterpart.

**Figure 6:**
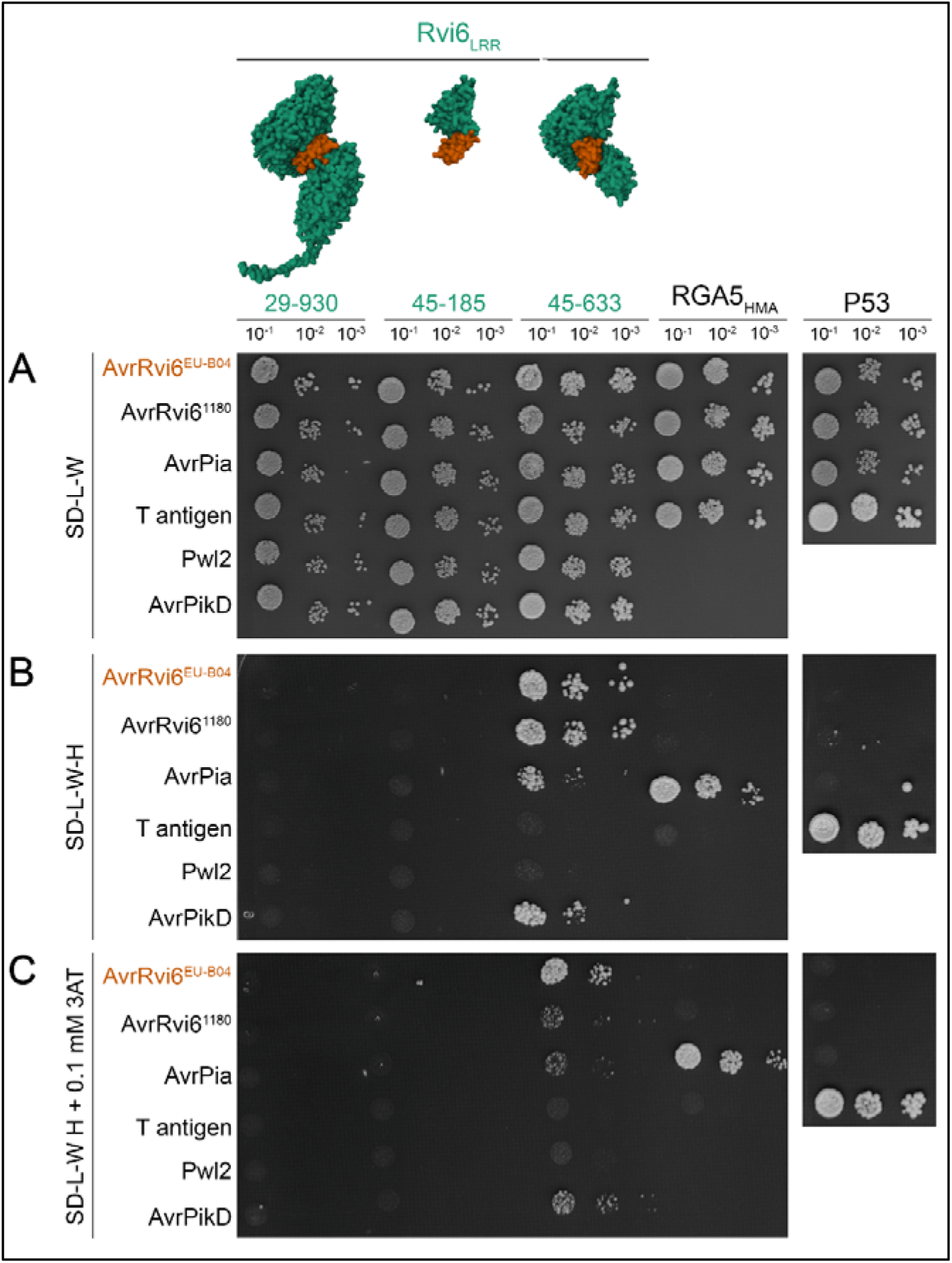
AvrRvi6 associates with Rvi6 in yeast. A) Schematic representation of Rvi6 extracellular LRR domain truncations used as bait (GAL4-BD) and growth of diploid yeast co-expressing the indicated bait and prey (GAL4-AD) fusions on non-selective control medium (SD-L-W). B) Interaction assays on moderately selective medium lacking histidine (SD-L-W-H). Rvi6_LRR45–633_ interacts with AvrRvi6^EU-B04^, AvrRvi6^1180^, AvrPikD, and AvrPia, but not Pwl2. Interaction specificity is confirmed by the absence of binding to RGA5_HMA_ (positive control for AvrPia; (Cesari et al. 2013) and murine p53 (negative control; (Li and Fields 1993). C) Stringent interaction assays on selective medium supplemented with 0.1 mM 3-amino-1,2,4-triazole (SD-L-W-H + 0.1 mM 3-AT). Robust growth is retained for AvrRvi6^EU-B04^, whereas the virulent variant AvrRvi6^1180^, AvrPikD, and AvrPia show weak growth. Serial 10-fold dilutions are shown from left to right. The experiment was repeated 3 times with similar results.

### 6. Mutations in predicted interface residues disrupt Rvi6-mediated AvrRvi6 recognition

Structural modeling of the Rvi6 – AvrRvi6 heterodimer predicted candidate interface residues on both partners (Figure 5), and Y2H assays confirmed their association in yeast (Figure 6). To functionally validate whether these specific contact residues govern receptor – effector recognition *in planta*, we mutated or deleted predicted interaction sites and evaluated their ability to trigger a hypersensitive response (HR) in *N. benthamiana*. We first grouped the predicted AvrRvi6 residues involved in Rvi6 recognition into four putative Rvi6-recognition sites (RS1–RS4; Figure 7A and 7B). We then engineered a series of alanine-substitution variants targeting these sites in AvrRvi6^EU-B04^: AvrRvi6^ΔRS-1234^ (all four sites mutated to Ala), AvrRvi6^ΔRS-134^ (sites 1, 3, and 4 mutated to Ala), AvrRvi6^ΔRS2^ (site 2 mutated to Ala), and AvrRvi6^ΔRS3^ (site 3 mutated to Ala). The wild-type AvrRvi6^EU-B04^ and the unrecognized virulent variant AvrRvi6^1180^ were included as positive and negative controls, respectively. Transient co-expression with Rvi6 revealed that AvrRvi6^ΔRS-1234^, AvrRvi6^ΔRS-134^ (Figure 7C), and AvrRvi6^ΔRS3^ (Figure 7D) completely failed to elicit an HR. In contrast, AvrRvi6^ΔRS2^ induced an intermediate phenotype, displaying significantly attenuated HR intensity compared to wild-type AvrRvi6^EU-B04^, yet significantly higher than the AvrRvi6^1180^ negative control (Figure 7D). Immunoblot analysis confirmed steady-state expression for all AvrRvi6 RS mutants in *N. benthamiana* leaves (Figure S6). Notably, AvrRvi6^ΔRS2^ exhibited a higher molecular weight shift on western blots, suggesting the accidental creation of a *de novo* N-glycosylation site near the C-terminal 6xHA epitope tag (Figure S6B). To test this hypothesis, we generated the non-glycosylated substitution AvrRvi6^N93A^; this variant restored the expected band size on immunoblots (Figure S6C) and triggered robust Rvi6-mediated HR (Figure 7D), confirming that the attenuation in AvrRvi6^ΔRS2^ was specifically attributable to the interface mutation rather than aberrant post-translational modification. To evaluate the reciprocal binding interface on the receptor, we deleted a solvent-exposed loop of Rvi6 (N91 to F103) involving clustered predicted contact residues, generating Rvi6Δ. Transient co-expression of Rvi6Δ with wild AvrRvi6^EU-B04^ completely abolished HR induction (Figure 7D). Together, these reciprocal mutational analyses demonstrate that the predicted contact residues on AvrRvi6 and an interacting loop region of Rvi6 are essential for receptor activation, providing functional validation of the structural binding interface predicted by AlphaFold-Multimer.

**Figure 7.**
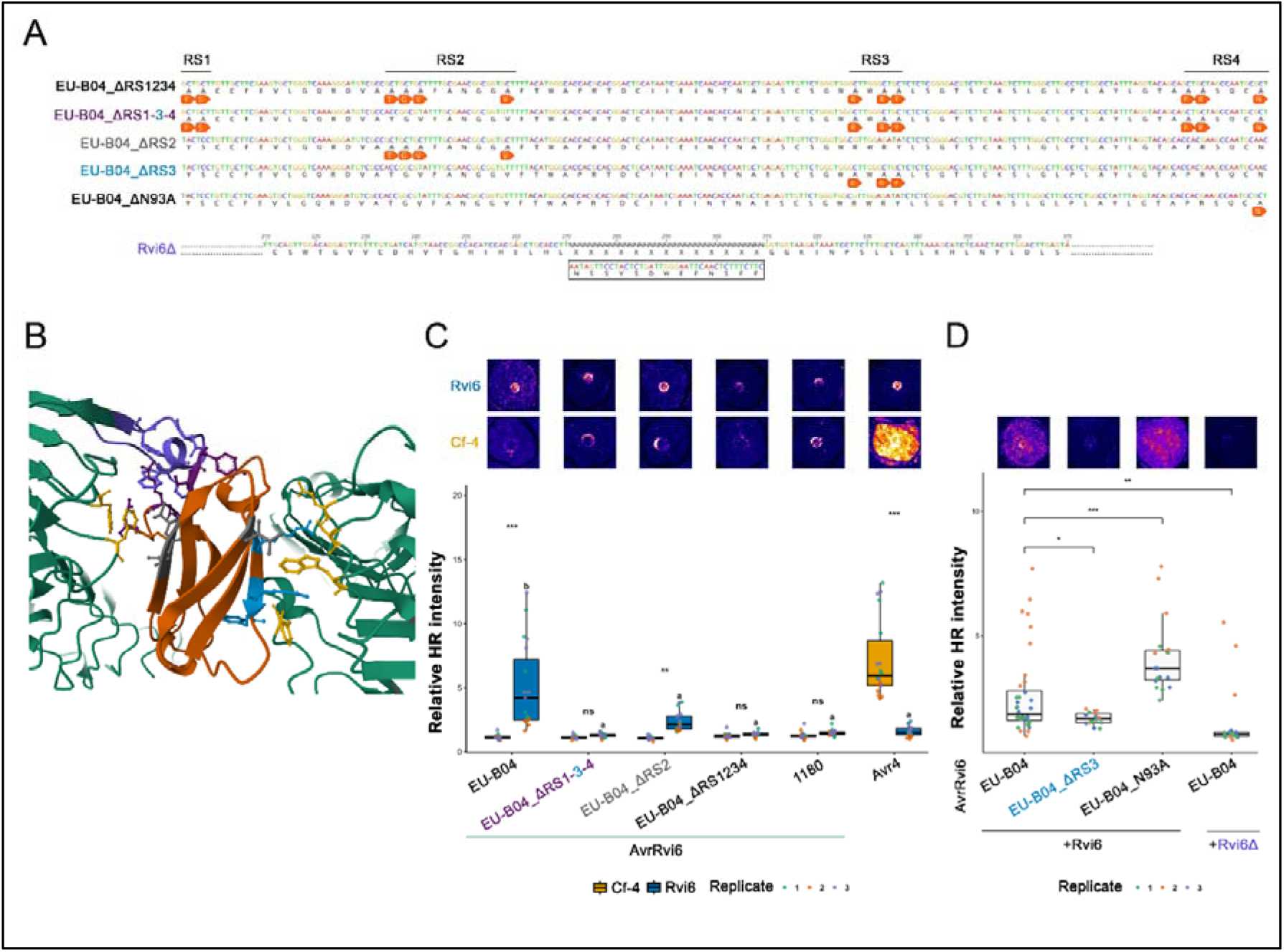
Mutations in predicted interface residues disrupt Rvi6-mediated AvrRvi6 recognition. A) DNA and protein sequences of expressed AvrRvi6 and Rvi6 generated mutants. Rvi6Δ in light purple and AvrRvi6 color-coded by mutated recognition sites: dark purple for RS1 and 4, grey for RS2 and blue for RS3. B) Schematic representation of predicted recognition sites on AvrRvi6 (sites RS1–RS4 color-coded as in (A)) and the complementary interaction residues on Rvi6 (all in yellow but the deleted loop in light purple). (C, D) Functional assessment of HR induction in *N. benthamiana* leaves upon transient co-expression of Rvi6 (in blue) and Cf-4 (in yellow) with AvrRvi6 alanine-substitution variants (C: AvrRvi6^ΔRS-1234^, AvrRvi6^ΔRS-134^; D: AvrRvi6^ΔRS2,^ AvrRvi6^ΔRS3^, AvrRvi6^N93A^) or the receptor loop mutant Rvi6Δ co-expressed with AvrRvi6^EU-B04^ (D). Wild-type AvrRvi6^EU-B04^ and the virulent variant AvrRvi6^1180^ serve as positive and negative controls, respectively. Points represent individual spots color-coded by replicate 3 independent biological replicates comprising 6 individual measures each were performed. All HR intensity signals are relative to non-infiltrated-leaf signal intensity. * p < 0.05, p < 0.01, *** p < 0.001, ns (not significant).

## Discussion

Several apple immune receptors conferring resistance to apple scab have been mapped or cloned, including *Rvi12*, *Rvi15*, and *Rvi6* (Galli et al. 2010; Schouten et al. 2014; Švara et al. 2024; Vinatzer et al. 2001). However, Rvi6 and AvrRvi6 currently represent the only pair for which both the host immune receptor and its cognate pathogen effector have been functionally characterized. The apoplastic perception of AvrRvi6 is coherent with the subcuticular lifestyle of *V. inaequalis*, which predominantly proliferates within the extracellular space between the cuticle and epidermal cell wall of apple tissues. Notably, AvrRvi6 is the first fungal MAX (Magnaporthe Avr and ToxB-like) effector shown to act in the apoplast (Sannier et al. 2025), raising key questions regarding how this structurally conserved effector family which was only described as recognized by intracellular nucleotide-binding leucine-rich repeat receptors (NLRs) so far is monitored outside host cells. In this study, we combined structural modeling, heterologous expression, live-cell imaging, and mutational analyses to dissect the molecular mechanisms governing Rvi6 – AvrRvi6 recognition, and we uncovered how naturally occurring virulent AvrRvi6 variants evade *Rvi6*-mediated immunity.

### LRR-RLP signaling paradigm is conserved for Rvi6 in the perennial crop *M. domestica*

Leucine-rich repeat receptor-like proteins (LRR-RLPs) lack an intracellular kinase domain and usually depend on other receptor-like kinases (RLKs) to transduce ligand perception across the plasma membrane (Liebrand et al. 2013; Postma et al. 2016). We demonstrated that *Rvi6*-mediated hypersensitive response (HR) in *N. benthamiana* strictly requires both the adaptor kinase *SOBIR1* and the co-receptor *BAK1*. This finding suggests that the central LRR-RLP signaling module established in model herbaceous annuals such as tomato Cf-4, Cf-9 and *Arabidopsis* RLP23 (Albert et al. 2015; Liebrand et al. 2013; Postma et al. 2016) is conserved in the woody perennial *Malus domestica.* In wild type plants, transient expression of the Rvi6–AvrRvi6^EU-B04^ pair elicited stronger HR in the absence of exogenous *MdSOBIR1* than in its presence likely because overexpressed *MdSOBIR1* competes with endogenous *NbSOBIR1*, which may couple more efficiently to downstream signaling components. In *bak1* knockout plants, the HR elicited by the control Cf-4 – Avr4 pair was not significantly reduced overall compared to WT plants, in contrast to the marked attenuation observed for Rvi6 – AvrRvi6^EU-B04^. Nevertheless, qualitative differences in cell death patterning were evident: expression of Cf-4 – Avr4 in *bak1* mutants resulted in patchy, confluent necrosis rather than uniform lesions covering the entire infiltration spot. This phenotypic difference likely reflects the strong HR elicited by the Cf-4 – Avr4 pair in *N. benthamiana*, together with functional redundancy among endogenous BAK1-like somatic embryogenesis receptor kinases (SERKs) that were not targeted during mutagenesis (Zönnchen et al. 2022). In contrast, both independent *Nbsobir1* mutant lines completely abolished HR triggered by Rvi6 and Cf-4, while functional complementation of *Nbsobir1* by apple *MdSOBIR1* fully restored HR induction. Consistent with this functional restoration at the cell surface, MdSOBIR1 localizes predominantly to the plasma membrane, placing it in the same subcellular compartment as the transmembrane receptor Rvi6. These results support a hypothesis of a model in which MdSOBIR1 constitutively interacts with Rvi6 at the plasma membrane, while MdBAK1 is recruited upon AvrRvi6 perception to form a competent tripartite complex that initiates downstream transphosphorylation and immune signaling. This assembly mechanism directly mirrors the ligand-induced activation paradigm established for the *Arabidopsis thaliana* LRR-RLP RLP23, which forms a constitutive pre-activation complex with SOBIR1 and recruits BAK1 only upon binding its cognate necrosis and ethylene-inducing peptide 1-like protein (NLP) ligand, such as nlp20 (Albert et al. 2015). Similarly, in tomato (*Solanum lycopersicum*), the LRR-RLPs Cf-4 and SlEIX2 constitutively pre-assemble with SlSOBIR1 at the plasma membrane, whereas the recruitment of SERK family co-receptors like SlBAK1/SERK3 is strictly ligand-dependent upon recognition of Avr4 and ethylene-inducing xylanase (EIX), respectively (Bar et al. 2010; Liebrand et al. 2013; Postma et al. 2016).

### The MAX-fold effector AvrRvi6 associates with Rvi6 at the plasma membrane

Recognition of pathogen-derived proteins by cell-surface plant immune receptors can rely on a direct physical interaction between the immune receptor and the pathogen protein. For example, the tomato RLP LeEIX2 directly binds the fungal elicitor EIX from *Trichoderma viride* (Ron and Avni 2004), while the *Arabidopsis* RLP RBPG1 was shown to physically associate with fungal endopolygalacturonases (Zhang et al. 2014). More recently, TurboID-based proximity labeling revealed associations between apoplastic effectors and their cognate receptors, including the biotinylation of the tomato receptor Cf-4 by Avr4 from *Fulvia fulva* and of Cf-2 by Avr2 from the same pathogen (Cebrailoglu et al. 2026). Importantly, although several RLPs recognize apoplastic fungal effectors, such as Ve1 – Ave1, direct physical interaction between the receptor and effector has not been demonstrated in these systems. Thus, direct perception of fungal effectors by RLPs remains poorly documented, highlighting the need for further investigation of the molecular mechanisms underlying RLP-mediated recognition of apoplastic effectors. Structural modeling and topological alignments revealed that AvrRvi6 shares core structural homology with the MAX effector family (Sannier et al. 2025). Previously known MAX effectors (e.g., AvrPikD and AvrPia) are translocated into the host cytoplasm and detected by intracellular NLRs via integrated heavy metal-associated (HMA) domains (Cesari et al. 2013; Maqbool et al. 2015; Ortiz et al. 2017). We previously demonstrated that the *V. inaequalis* MAX effector is an effector with apoplastic activities (Sannier et al. 2025). Our results showed a main localization of the LRR-RLP Rvi6 at the plasma membrane, which is coherent with its features of transmembrane LRR-RLP protein and the previously published localization data (Chen et al. 2025; Vinatzer et al. 2001). Our data demonstrate that the presence of Rvi6 relocalizes AvrRvi6 to the cell-surface. Our yeast two-hybrid (Y2H) assays show that the central extracellular LRR domain of Rvi6 (residues 45–633) associates with AvrRvi6. This association mirrors the apoplastic perception mechanism of tomato Cf-4 toward the chitin-binding effector Avr4 (Cebrailoglu et al. 2026; Postma et al. 2016), as well as the *N. benthamiana* LRR-RLP RXEG1, which directly binds the oomycete glycoside hydrolase XEG1 to recruit BAK1 (Sun et al. 2022). Multiple AvrRvi6 alleles are recognized by Rvi6, while Rvi6LRR_45–633_ also exhibited weak interaction signals in yeast with some other MAX effectors, although robust yeast growth was retained exclusively for the avirulent allele AvrRvi6^EU-B04^. These observations suggest that the Rvi6 LRR ectodomain may display an intrinsic structural predisposition toward the MAX scaffold, whereas effective immune activation requires stronger, sequence-specific contacts restricted to cognate avirulent alleles. Nevertheless, Rvi6 appears to possess a relatively broad recognition capacity, reminiscent of RGA5, whose integrated HMA domain directly binds the *M. oryzae* effectors AVR-Pia and AVR1-CO39 (Cesari et al. 2013; Guo et al. 2018). Several studies have demonstrated the potential of exploiting such recognition properties through protein engineering. Structure-guided engineering of the Pikp–HMA domain in rice showed that a single interface mutation could enhance binding affinity to previously unrecognized AVR-Pik variants, thereby expanding the receptor’s effector recognition spectrum and cell-death response (De la Concepcion et al. 2019). Similarly, the molecular mimicry mechanism underlying MLA3 recognition was exploited to engineer a chimeric SR50 receptor in barley by transferring the effector-binding interface of MLA3 into SR50 (Gómez De La Cruz et al. 2026). This resulted in a receptor with dual recognition activities that conferred resistance to two major cereal pathogens in transgenic barley lines. Together, these findings highlight the potential of rational receptor engineering to expand immune recognition and confer resistance to multiple pathogens. In this context, the broad recognition capacity of Rvi6 makes it a promising starting point for engineering an immune receptor with an expanded recognition spectrum against apoplastic MAX effectors.

### Virulent AvrRvi6 alleles escape Rvi6 recognition by avoiding apoplastic accumulation

At the sequence level, three AvrRvi6 single nucleotide polymorphisms were previously associated with virulence towards *Rvi6*-carrying apple genotypes (Sannier et al. 2025). Our structural analyses indicate that these polymorphisms do not generally result in major alterations of the AvrRvi6 fold. The virulent AvrRvi6^2917^ allele, which clusters with avirulent alleles, retains the same overall backbone architecture, suggesting that its virulence is unlikely to result from a large-scale structural rearrangement. Instead, its C21S substitution may impair the formation of a disulfide bond. Disulfide bonds therefore appear to be important for maintaining the structural and/or functional properties required for *Rvi6* recognition. The AvrRvi6^1180^ and AvrRvi6^1153^ alleles were more structurally divergent, although their overall similarity to avirulent alleles remained high (RMSD < 0.5). Both clustered as structural outliers relative to the main avirulent group, suggesting that their specific combinations of polymorphisms may subtly affect protein conformation. In AvrRvi6^1180^, F36S is particularly notable because it represents the only virulence-associated substitution predicted to be exposed rather than buried. Replacement of phenylalanine by serine at this position could disrupt local hydrophobic packing and introduce a polar group into a partially buried environment, potentially reducing local structural stability (Eriksson et al. 1992). Alternatively, the effect of F36S may depend on its combination with the other polymorphisms present in AvrRvi6^1180^, collectively shifting the protein away from the structural configuration observed in avirulent alleles. AvrRvi6^1153^ was the most divergent allele relative to AvrRvi6^EU-B04^, which may similarly result from the combined effect of its polymorphisms, including C50R, which disrupts a predicted disulfide bond. Thus, although individual virulence-associated substitutions may affect local structural features, the structural differences identified here do not provide a straightforward explanation for the loss of *Rvi6* recognition. Instead, analysis of subcellular localization provided a more compelling explanation for the virulent phenotype. In *N. benthamiana*, the two virulent alleles tested, AvrRvi6^1180^ and AvrRvi6^1153^, failed to accumulate in the apoplast to the same extent as the avirulent AvrRvi6^EU-B04^ allele. Their fluorescence was instead predominantly detected in the vacuole. Because all three constructs were expressed with the same PR1α secretion signal and no polymorphism has been detected in the endogenous AvrRvi6 signal peptide among *V. inaequalis* isolates (Sannier et al. 2025), these differences are unlikely to result from variation in the secretion signal itself. The observed localization may therefore reflect either inefficient secretion or reinternalization of the virulent proteins. One possible explanation is that structural or biochemical properties conferred by their polymorphisms reduce their stability, leading to their removal through intracellular quality-control pathways. In particular, chaperone-assisted selective autophagy could contribute to the trafficking of unstable proteins towards the vacuole for degradation (Marshall and Vierstra 2018). Another possible explanation is that AvrRvi6 could be hijacking the plant Vacuolar Sorting Receptor-mediated trafficking to evade host immune detection. Indeed, AvrRvi6 may be associating with a Vacuolar Sorting Receptor (VSR) to be directed to the prevacuolar compartment (PVC) and then the vacuole such as it was demonstrated for the *S. scitamineum* effector *SsPE15* (Ling et al. 2025). If a similar localization defect occurs during *V. inaequalis* infection, AvrRvi6^1180^ and AvrRvi6^1153^ might consequently escape *Rvi6* surveillance by failing to accumulate in the apoplast, where recognition is expected to occur. Their altered localization would, however, raise an important evolutionary trade-off. If these alleles are inefficiently secreted or subsequently retained within fungal or host cells, they may no longer efficiently perform the functions normally associated with AvrRvi6 during infection. In this respect, AvrRvi6^1180^ and AvrRvi6^1153^ may represent an intermediate stage towards pseudogenization, similar to the loss-of-function mechanisms previously proposed for strains carrying transposon insertions associated with the avirulent AvrRvi6^0301^ or the truncated AvrRvi6^2828^ allele (Sannier et al., 2025). Such a mechanism would be consistent with strong selection imposed by the broad recognition capacity of *Rvi6*, favoring the loss or sequestration of the effector rather than extensive modification of its sequence.

### Rvi6 recognizes AvrRvi6 through multiple interaction surfaces

Our AlphaFold-based modelling of the Rvi6–AvrRvi6 complex identified several candidate interaction sites on both proteins. Functional analysis of these residues provided experimental support for the predicted interaction interface: mutation of all candidate sites in AvrRvi6, as well as of most of the corresponding sites in Rvi6, impaired or abolished recognition. These results support an association between the two proteins and indicate that recognition depends on multiple regions of the complex rather than on a single dominant interaction site. In particular, mutation of RS2 only partially reduced recognition, whereas the RS3 mutation or combinations of mutations resulted in more drastic loss of HR. This suggests that several contact regions may contribute cooperatively to the interaction, allowing recognition to be maintained despite changes at individual residues. The localization of the predicted Rvi6 interaction sites within mostly exposed loops is consistent with previous studies showing that extracellular recognition by plant receptor-like proteins can involve structurally flexible regions, including inter-LRR loops (van der Hoorn et al. 2005). These regions are subject to fewer structural constraints than the conserved LRR core and can therefore tolerate sequence variation while preserving the overall receptor fold. Such an architecture could provide Rvi6 with both evolutionary flexibility and a broad recognition surface. Extended or flexible loops may form three-dimensional binding pockets or adaptable surfaces capable of accommodating structurally diverse effector molecules. This may help explain why Rvi6 retains the capacity to recognize a large proportion of the AvrRvi6 sequence diversity identified in natural populations despite the accumulation of numerous SNPs.

### Concluding remarks

Taken together, our results suggest that Rvi6 – AvrRvi6 co-evolution may not primarily follow a classical sequence-based arms race in which the effector progressively accumulates mutations directly within the recognition interface. Rather, the broad recognition spectrum of *Rvi6*, the limited number of virulence-associated AvrRvi6 polymorphisms, the apparent ability of Rvi6 to associate with a virulent allele under certain experimental conditions, and the altered intracellular localization of AvrRvi6^1180^ and AvrRvi6^1153^ collectively support an alternative mechanism of adaptation. In this model, selection imposed by *Rvi6* favors changes that prevent AvrRvi6 from reaching or persisting in the apoplast, thereby reducing its exposure to the receptor, rather than extensive modification of the effector surface itself. Determining the localization and secretion behavior of these alleles in apple leaves during *V. inaequalis* infection will be critical to test this hypothesis. More broadly, these findings suggest that the stability of both the receptor and effector recognition machinery may be maintained during host–pathogen co-evolution, with adaptation occurring through changes in protein trafficking and cellular compartmentalization rather than through continuous diversification of the recognition interface.

## Materials and methods

### Plant Materials and Growth Conditions

*Nicotiana benthamiana* wild-type (WT) plants, alongside the previously described mutant lines *Nbnrc234_5.5.1* (Wu et al. 2020)*, Nbbak1* (Zönnchen et al. 2022), *Nbsobir1-1* and *Nbsobir1-2* (Huang et al. 2021), were grown under controlled greenhouse conditions at approximately 20°C under a 16 h light / 8 h dark photoperiod. Five- to seven-week-old plants were used for all transient agroinfiltration, live-cell imaging, and biochemical experiments.

### Plasmid Construction and Site-Directed Mutagenesis

Binary expression vectors were modified using the Modular Cloning (MoClo) system to insert target genes (Supplemental table 1) for the transformation of *Agrobacterium tumefaciens* strains GV3101 or GV3103. Selection markers were chosen based on the specific plasmid and host bacterial strain resistances (Supplemental table 1). To ensure proper recognition by the apple receptor Rvi6, *AvrRvi6* alleles were cloned downstream of the *Nicotiana tabacum* PR1α signal peptide sequence to direct apoplastic secretion (Van der Hoorn et al. 2000). For protein expression validation via western blot, a 6xHA epitope tag was fused to the C-terminus of each gene. Fluorescent constructs were assembled utilizing the Golden Gate MoClo Toolkit (plant parts I and II kits; Engler et al. 2014; Gantner et al. 2018; Weber et al. 2011). In silico cloning design and sequence alignments were performed using Geneious Prime (v9.0.5). All plasmid constructs were verified by PCR and Sanger sequencing before *Agrobacterium* transformation. Detailed plasmids construction schemes are provided in Supplemental Table 1. Effector constructs were tagged at their C-termini with either 6xHA or mCherry. For subcellular localization, *Rvi6* was fused to C-terminal mEGFP, mVenus, or mStayGold reporters. The plasma membrane-localized RXLR effector PITG16242 was used as a cell membrane marker, fused to mCherry or GFP (Petre et al. 2021). Coding sequences for *Rvi6* and its extracellular domain truncations—*Rvi6_LRR29–930_*, *Rvi6_LRR45–633_*, and *Rvi6_LRR45–185_*—were amplified from existing expression vectors. The *MdSOBIR1* coding sequence was amplified from *M. domestica* cDNA. Alanine substitutions targeting predicted Rvi6 recognition sites (RS1–RS4) in AvrRvi6^EU-B04^ (*AvrRvi6^ΔRS-1234^*, *AvrRvi6^ΔRS-134^*, *AvrRvi6^ΔRS2^*, and *AvrRvi6^ΔRS3^*) and the deletion targeting a predicted effector-binding loop of Rvi6 (*Rvi6Δ*), were synthetized by Twist Bioscience (South San Francisco, CA, USA). The de-glycosylation mutant *AvrRvi6^N93A^* was generated using the Q5® Site-Directed Mutagenesis Kit (New England Biolabs, Ipswich, MA, USA), according to the manufacturer’s instructions. Heterologous effectors from *Magnaporthe oryzae* (*AvrPikD*, *AvrPia*, and *Pwl2*) lacking native secretion signals were fused downstream of PR1α^SP^.

### Agrobacterium tumefaciens-Mediated Transient Transformation of N. benthamiana

For transient expression in *Nicotiana benthamiana*, *Agrobacterium tumefaciens* strains harboring the desired constructs were initially propagated at 28°C on antibiotic-supplemented LB agar for 48 hours (see Supplemental Table 1 for specific selection criteria). Prior to infiltration, bacterial cells were collected and conditioned by a 4-hour dark incubation in 2 mL of induction buffer containing 50 mM MES (pH 5.6) and 200 µM acetosyringone. The density of each induced suspension was calibrated to an optical density at 600 nm (OD_600_) of 2. To assess hypersensitive responses (HR) or perform confocal microscopy, individual cultures were pooled in equal ratios to yield final per-construct OD_600_ targets of 0.25 and 0.1, respectively. Using a needleless syringe, these mixtures were introduced into the abaxial sides of leaves from 5- to 7-week-old plants. Experimental robustness was maintained by infiltrating six distinct leaves distributed across three separate plants per treatment, with three independent biological replicates. Plants were maintained in the greenhouse at ∼20°C under a 16 h photoperiod for 3 to 4 days before phenotypic evaluation.

### Hypersensitive Response (HR) Assays and Phenotypic Quantification

Quantification of HR cell death was conducted 4 days post-infiltration using red-light imaging on a ChemiDoc™ MP Imaging System (Bio-Rad, Hercules, CA, USA) to record chlorophyll fluorescence quenching (Landeo Villanueva et al. 2021). Three biological replicates for each infiltration combination were performed on six different *N. benthamiana* leaves on two different plants each time. Median pixel intensity within each infiltration spot was quantified using FIJI (Schindelin et al. 2012) and normalized against the non-infiltrated area of each leaf. All statistical analyses and graphical representations were performed in R (version 2025.09.2; R Core Team, 2025), utilizing the tidyverse, lme4, lmerTest, emmeans, and multcomp packages. Depending on the assay and experimental design, response variables were analyzed using either non-parametric tests, parametric tests, or linear mixed-effects models (LMM) fitted by restricted maximum likelihood (REML), incorporating biological replicates and/or plant identity as random intercepts to control for batch-to-batch variation. For pairwise comparisons within the *bak1* dataset, data normality was verified using the Shapiro–Wilk test; normally distributed datasets were evaluated using two-tailed Student’s *t*-tests, whereas non-normally distributed data were assessed via Wilcoxon rank-sum tests. For *MdSOBIR1* complementation assays across genetic backgrounds (*sobir1-1, sobir1-2,* and wild-type), relative hypersensitive response (HR) intensity values were log-transformed (log(y + 1)). Data were then analyzed using an LMM modeling plant genotype, receptor–effector interaction, and MdSOBIR1 complementation as fixed interaction terms, with biological replicate and individual plant ID included as random intercepts. Post hoc comparisons within each genotype and interaction group were conducted via estimated marginal means (emmeans) with back-transformed response ratios. For functional assays evaluating AvrRvi6 and Rvi6 mutants in *N. benthamiana*, quantitative pixel intensity measurements (median values) were analyzed using an LMM with experimental replicate specified as a random intercept. Post hoc pairwise contrasts against the reference pair (Rvi6-AvrRvi6^EU-B04^) were conducted using Dunnett’s test, while broader multi-condition contrasts were adjusted using Tukey’s Honest Significant Difference (HSD). For all tests, statistical significance was defined at *p* < 0.05 (* *p* < 0.05, \*\**p* < 0.01, *** *p* < 0.001; ns, not significant).

### Laser-Scanning Confocal Microscopy

*N. benthamiana* leaf fragments transiently expressing fluorescent fusion proteins were sampled at 3 days post-infiltration (dpi), vacuum-infiltrated with water, and mounted between glass slides and coverslips. Imaging was performed on a Nikon AX laser-scanning confocal microscope (IMAC platform, SFR QuaSaV, Angers) equipped with a Plan Fluor 40x oil-immersion objective (NA 1.30, DIC H N2). Fluorophores were excited and detected sequentially to prevent spectral bleed-through: mEGFP/GFP/mStayGold (excitation 488 nm, emission 499–533 nm), mVenus (excitation 514 nm, emission 525–555 nm), mCherry (excitation 561 nm, emission 592–625 nm), and chlorophyll autofluorescence (excitation 640 nm, emission 662–737 nm). All images were processed using FIJI (Schindelin et al. 2012). Line-profile fluorescence intensity plots were generated using the Plot Profile tool and pixel values were normalized to the maximum fluorescence intensity per channel. Relative fluorescence intensity plots were plotted using Microsoft Excel (v365; Microsoft).

### Western Blot Analysis

Apoplastic wash fluids (AF) were collected from agroinfiltrated *N. benthamiana* leaves at 3 dpi. Excised leaves were submerged in ice-cold water and vacuum-infiltrated for 10 min. Leaf surfaces were blotted dry with absorbent paper, rolled, and inserted into empty 20 mL needleless syringes nested within 50 mL conical tubes. AFs were collected by centrifugation at 2000g for 10 min at 4°C and immediately flash-frozen in liquid nitrogen.

For total crude protein extraction, 1.5 cm² leaf discs from agroinfiltrated tissue were ground in liquid nitrogen. AF and crude protein extracts were supplemented with Laemmli buffer and denatured at 95°C for 5 min. Proteins were separated on 12% SDS-PAGE gels and electroblotted onto membranes. Target proteins were probed using primary antibodies: high-affinity rat monoclonal anti-HA (Roche, 12013819001; 1:2000) or rabbit polyclonal anti-RFP (Rockland, 600-401-379; 1:1000) for mCherry fusions, followed by chemiluminescent detection.

To confirm the steady-state expression of bait and prey fusion proteins in the Y2H experiment, total protein was extracted from yeast strains cultured in liquid SD medium using the Kushnirov post-alkaline extraction protocol (Kushnirov 2000). For immunoblotting, equal volumes of yeast lysates were resolved on 4-20% gradient SDS-PAGE gels and electroblotted onto PVDF membranes. Membranes were blocked for 1 hour at room temperature in 5% (w/v) non-fat dry milk prepared in TBS-T (50 mM Tris-HCl pH 7.5, 150 mM NaCl, 0.1% [v/v] Tween-20). Bait fusions were detected using a mouse monoclonal anti-GAL4-BD primary antibody, while prey fusions were probed with a horseradish peroxidase (HRP)-conjugated anti-HA primary antibody. After three washes in TBS-T, Bait fusion membranes were incubated with HRP-conjugated goat anti-mouse IgG secondary antibody for 1 hour at room temperature. Protein bands were visualized using enhanced chemiluminescence (ECL) substrate on a ChemiDoc™ MP Imaging System (Bio-Rad, Hercules, CA, USA). Total protein loading was verified by Ponceau S staining prior to blocking.

### Yeast Two-Hybrid (Y2H) Assays

Y2H assays were carried out using the Matchmaker Gold Yeast Two-Hybrid System (Clontech). Mature effector variants lacking signal peptides and the control T antigen were cloned into pGADT7 (GAL4 activation domain; AD prey), and Rvi6 extracellular LRR truncations (Rvi6_LRR29–930_, Rvi6_LRR45–633_ and Rvi6_LRR45–633_), the RGA5_HMA_, or control p53 were cloned into pGBKT7 (GAL4 DNA-binding domain; BD bait). The bait constructs were transformed into the *Saccharomyces cerevisiae* strain Gold, while prey constructs were transformed into strain Y187. Haploid transformants were isolated on selective synthetic defined (SD) agar media lacking tryptophan (SD–Trp) for bait strains or leucine (SD–Leu) for prey strains. For mating, single colonies of corresponding bait and prey strains were inoculated into 2x YPDA liquid medium and co-cultured at 30°C with gentle agitation (120 rpm) for 24 hours to facilitate zygote formation. Aliquots of the mating cultures were subsequently plated onto double-dropout medium lacking leucine and tryptophan (SD-Leu-Trp; SD-L-W) to select for surviving diploid cells containing both prey and bait expressing plasmids. Protein-protein interactions were screened on selective triple-dropout medium lacking leucine, tryptophan, and histidine (SD-Leu-Trp-His; SD-L-W-H). Stringent interaction assays were assessed on SD-L-W-H medium supplemented with 0.1 and mM 3-amino-1,2,4-triazole (3-AT). Ten-fold serial dilutions (OD600 = 1.0, 0.1, 0.01) were spotted onto agar plates and incubated at 30°C for 3 to 5 days. Auto-activation and non-specific backgrounds were controlled by mating all target lines against strains harboring negative controls (P53 for effectors and T antigen for receptors). A minimum of three biological repeats were performed for each association test.

### In Silico Structural Modeling and Interface Analysis

Monomer tertiary structures for all 20 natural *V. inaequalis* AvrRvi6 alleles were modeled using AlphaFold2 (Jumper et al. 2021). Heterodimeric receptor–effector complexes between Rvi6 and AvrRvi6 variants were predicted using AlphaFold-Multimer (v2.3 ; Evans et al. 2021). Complex model confidence was determined using the weighted interface metric: 0.8 ipTM + 0.2pTM (Evans et al. 2021). Pairwise structural alignments and root-mean-square deviation (RMSD) calculations were computed in UCSF ChimeraX (Pettersen et al. 2021). Structural distance dendrograms were constructed using DALI (Holm 2020). Solvent-accessible surface area (SASA) for natural polymorphic sites was calculated using standard probe radii (1.4 Å) to differentiate core-buried residues from surface-accessible positions on PDB viewer (Sehnal et al. 2021).

## Acknowledgements

We thank Isabel Guerrero-Zepeda for assistance with some of the experimental procedures. We thank Stella Césari for providing the RGA5, AvrPikD, Pwl2 and AvrPia plasmids and templates; Benjamin Petre for providing the GFP-PITG16242 and PITG16242-mCherry plasmids; Sophien Kamoun for providing the *Nbnrc234* mutants; Johannes Stuttmann and Matthieu Joosten for the *sobir1* and *bak1 N. benthamiana* mutants. We thank the IMAC (Cellular Imaging) technical platform of SFR 4207 QUASAV, Université d’Angers, especially Aurélia Rolland, for providing access to confocal microscopy equipment and technical guidance. The Angers Plant Phenotyping Facility PHENOTIC (https://doi.org/10.17180/ykbz-2v85) is acknowledged for plant production in the glasshouses.

## Author contributions

Conceptualizations: SB, BLC, MB.

Methodology: SB, MB

Investigation: SB, MS, MB

Funding acquisition: BLC, MB

Writing - original draft: SB, MB

Writing – review & editing: SB, MS, BLC, MB

## Figure legends

<u>Figure S1: MdSOBIR1 is localized at the plasma membrane in *N. benthamiana.*</u> Subcellular localization of MdSOBIR1, fused to an mVenus fluorescent tag, in *N. benthamiana* leaf epidermal cells at 3 days post-infiltration (dpi) analyzed by laser-scanning confocal microscopy. MdSOBIR1 was co-expressed with the apoplastic avirulent AvrRvi6^EU-B04^ carrying the plant PR1α signal peptide and a C-terminal mCherry tag (PR1αSP-AvrRvi6ΔSP-mCherry magenta/red), both under a double 35S promoter. MdSOBIR1 localizes to the membrane. The avirulent variant EU-B04 localizes predominantly to the extracellular space (apoplast). Scale bars =10 µm. Signal intensity from point “a” to “b” is shown in a plot. Experiment was repeated at least twice.

<u>Figure S2: AvrRvi6 alleles sequence alignment</u>. All AvrRvi6 alleles studied, except for the truncated AvrRvi6^2828^ allele have been aligned. polymorphisms present in both virulent and avirulent alleles are highlighted in yellow, those present only in virulent alleles are shown in red. Virulent allele names are written in red, avirulent ones in black.

<u>Figure S3:</u> <u>Structural dendrogram of AvrRvi6 alleles</u>. All AvrRvi6 alleles studied, except for the truncated AvrRvi6^2828^ allele have been 3D modeled and structurally aligned. Avirulent allele names are written in green and virulent ones in red.

<u>Figure S4:</u> <u>Confidence score analysis of Rvi6-AvrRvi6 complexes</u>. Complex model confidence was determined using the weighted interface metric: 0.8 ipTM + 0.2pTM (Evans et al. 2021). Virulent allele names are written in red, avirulent ones in green.

<u>Figure S5: Rvi6_LRR45–633_ associates with AvrRvi6 and some other MAX effectors in yeast but only AvrRvi6 is recognized triggering an HR in *N. benthamiana.*</u> A) Immunoblot analysis confirming the steady-state expression of proteins involved in the Y2H experiment (Figure 6). B) Structural alignment of MAX effectors tested in Y2H experiment with the reference AvrRvi6^EU-B04^. C) Functional assessment of HR induction in *N. benthamiana* leaves upon transient co-expression of Rvi6 with different MAX effectors. D) Immunoblot analysis confirming the steady-state expression of effectors expressed in *N. benthamiana* in (D).

<u>Figure S6:</u> <u>AvrRvi6 recognition-site-mutants are well expressed in *N. benthamiana.*</u> A) Immunoblot analysis confirming the steady-state expression of AvrRvi6^EU-B04^, AvrRvi6^1180^ and the AvrRvi6 mutants (AvrRvi6^ΔRS1234^, AvrRvi6^ΔRS134^ and AvrRvi6^ΔRS2^), in *N. benthamiana* during HR induction tests (Figure 7C). B) Protein sequence of AvrRvi6^ΔRS2ΔSP^-6xHA with indicated potential N-glycosylation motifs. C) Immunoblot analysis confirming the steady-state expression of AvrRvi6^ΔRS3^, AvrRvi6^N93A^, AvrRvi6^EU-B04^ in *N. benthamiana* during HR induction tests (Figure 7D) Avr4 is used as a positive control.

## References

Albert, I., Böhm, H., Albert, M., Feiler, C. E., Imkampe, J., Wallmeroth, N., Brancato, C., Raaymakers, T. M., Oome, S., Zhang, H., Krol, E., Grefen, C., Gust, A. A., Chai, J., Hedrich, R., van den Ackerveken, G., and Nürnberger, T. 2015. An RLP23–SOBIR1–BAK1 complex mediates NLP-triggered immunity. Nat. Plants 1:15140.

Bar, M., Sharfman, M., Ron, M., and Avni, A. 2010. BAK1 is required for the attenuation of ethylene-inducing xylanase (Eix)-induced defense responses by the decoy receptor LeEix1. Plant J. 63:791–800.

Belfanti, E., Silfverberg-Dilworth, E., Tartarini, S., Patocchi, A., Barbieri, M., Zhu, J., Vinatzer, B. A., Gianfranceschi, L., Gessler, C., and Sansavini, S. 2004. The *HcrVf2* gene from a wild apple confers scab resistance to a transgenic cultivated variety. Proc. Natl. Acad. Sci. U.S.A. 101:886–890.

Boevink, P. C., Wang, X., McLellan, H., He, Q., Naqvi, S., Armstrong, M. R., Zhang, W., Hein, I., Gilroy, E. M., Tian, Z., and Birch, P. R. J. 2016. A *Phytophthora infestans* RXLR effector targets plant PP1c isoforms that promote late blight disease. Nat. Commun. 7:10311.

Bowen, J. K., Mesarich, C. H., Bus, V. G. M., Beresford, R. M., Plummer, K. M., and Templeton, M. D. 2011. *Venturia inaequalis*: the causal agent of apple scab. Mol. Plant Pathol. 12:105–122.

Cebrailoglu, N., Budak, E., Villanueva, S. L., Schol, C. R., ter Waarbeek, C., Evertsz, K., Boeren, S., and Joosten, M. H. A. J. 2026. Effector-centred proximity-dependent labelling enables the discovery of cell-surface immune receptors in plants. bioRxiv. doi:10.1101/2026.03.23.713615.

Cesari, S., Thilliez, G., Ribot, C., Chalvon, V., Michel, C., Jauneau, A., Rivas, S., Alaux, L., Kanzaki, H., Okuyama, Y., Morel, J.-B., Fournier, E., Tharreau, D., Terauchi, R., and Kroj, T. 2013. The rice resistance protein pair RGA4/RGA5 recognizes the *Magnaporthe oryzae* effectors AVR-Pia and AVR1-CO39 by direct binding. Plant Cell 25:1463–1481.

Chen, A., Guo, H., Jiang, H., Yang, W., Wang, X., and Xu, L. 2025. Rvi6-mediated defense mechanisms against apple scab. BMC Plant Biol. 25:1137.

De Guillen, K., Ortiz-Vallejo, D., Gracy, J., Fournier, E., Kroj, T., and Padilla, A. 2015. Structure analysis uncovers a highly diverse but structurally conserved effector family in phytopathogenic fungi. PLoS Pathog. 11.

De la Concepcion, J. C., Franceschetti, M., MacLean, D., Terauchi, R., Kamoun, S., and Banfield, M. J. 2019. Protein engineering expands the effector recognition profile of a rice NLR immune receptor. eLife 8:e47713

Deng, C. H., Plummer, K. M., Jones, D. A. B., Mesarich, C. H., Shiller, J., Taranto, A. P., Robinson, A. J., Kastner, P., Hall, N. E., Templeton, M. D., and Bowen, J. K. 2017. Comparative analysis of the predicted secretomes of Rosaceae scab pathogens *Venturia inaequalis* and *V. pirina* reveals expanded effector families and putative determinants of host range. BMC Genomics 18:339.

Engler, C., Youles, M., Gruetzner, R., Ehnert, T.-M., Werner, S., Jones, J. D. G., Patron, N. J., and Marillonnet, S. 2014. A Golden Gate modular cloning toolbox for plants. ACS Synth. Biol. 3:839–843.

Eriksson, A. E., Baase, W. A., Zhang, X. J., Heinz, D. W., Blaber, M., Baldwin, E. P., and Matthews, B. W. 1992. Response of a protein structure to cavity-creating mutations and its relation to the hydrophobic effect. Science 255:178–183.

Evans, R., O’Neill, M., Pritzel, A., Antropova, N., Senior, A., Green, T., Žídek, A., Bates, R., Blackwell, S., Yim, J., Ronneberger, O., Bodenstein, S., Zielinski, M., Bridgland, A., Potapenko, A., Cowie, A., Tunyasuvunakool, K., Jain, R., Clancy, E., Kohli, P., Jumper, J., and Hassabis, D. 2021. Protein complex prediction with AlphaFold-Multimer. bioRxiv. doi:10.1101/2021.10.04.463034.

Galli, P., Patocchi, A., Broggini, G. A. L., and Gessler, C. 2010. The Rvi15 (Vr2) apple scab resistance locus contains three TIR-NBS-LRR genes. Mol. Plant-Microbe Interact. 23:608–617.

Gantner, J., Ordon, J., Ilse, T., Kretschmer, C., Gruetzner, R., Löfke, C., Dagdas, Y., Bürstenbinder, K., Marillonnet, S., and Stuttmann, J. 2018. Peripheral infrastructure vectors and an extended set of plant parts for the Modular Cloning system. PLoS ONE 13.

Gómez De La Cruz, D., Ingram, T., Zdrzałek, R., Taylor, J., Wawryk-Khamdavong, A., Bachowska, K., Banfield, M. J., Talbot, N. J., and Moscou, M. J. 2026. Molecular mimicry of a pathogen virulence target by a plant immune receptor. Science. 392:1050–1055

Guo, L., Cesari, S., de Guillen, K., Chalvon, V., Mammri, L., Ma, M., Meusnier, I., Bonnot, F., Padilla, A., Peng, Y.-L., Liu, J., and Kroj, T. 2018. Specific recognition of two MAX effectors by integrated HMA domains in plant immune receptors involves distinct binding surfaces. Proc. Natl. Acad. Sci. U.S.A. 115:11637–11642.

Gust, A. A., and Felix, G. 2014. Receptor-like proteins associate with SOBIR1-type of adaptors to form bimolecular receptor kinases. Curr. Opin. Plant Biol. 21:104–111.

Holm, L. 2020. DALI and the persistence of protein shape. Protein Sci. 29:128–140.

van der Hoorn, R. A. L., Wulff, B. B. H., Rivas, S., Durrant, M. C., van der Ploeg, A., de Wit, P. J. G. M., and Jones, J. D. G. 2005. Structure–function analysis of Cf-9, a receptor-like protein with extracytoplasmic leucine-rich repeats. Plant Cell 17:1000–1015.

Huang, W. R. H., Schol, C., Villanueva, S. L., Heidstra, R., and Joosten, M. H. A. J. 2021. Knocking out SOBIR1 in *Nicotiana benthamiana* abolishes functionality of transgenic receptor-like protein Cf-4. Plant Physiol. 185:290–294.

Jumper, J., Evans, R., Pritzel, A., Green, T., Figurnov, M., Ronneberger, O., Tunyasuvunakool, K., Bates, R., Žídek, A., Potapenko, A., Bridgland, A., Meyer, C., Kohl, S. A. A., Ballard, A. J., Cowie, A., Romera-Paredes, B., Nikolov, S., Jain, R., Adler, J., Back, T., Petersen, S., Reiman, D., Clancy, E., Zielinski, M., Steinegger, M., Pacholska, M., Berghammer, T., Bodenstein, S., Silver, D., Vinyals, O., Senior, A. W., Kavukcuoglu, K., Kohli, P., and Hassabis, D. 2021. Highly accurate protein structure prediction with AlphaFold. Nature 596:583–589.

Kruijt, M., Kip, D. J., Joosten, M. H. A. J., Brandwagt, B. F., and de Wit, P. J. G. M. 2005. The Cf-4 and Cf-9 resistance genes against *Cladosporium fulvum* are conserved in wild tomato species. Mol. Plant-Microbe Interact. 18:1011–1021.

Kushnirov, V. V. 2000. Rapid and reliable protein extraction from yeast. Yeast 16:857–860.

Landeo Villanueva, S., Malvestiti, M. C., van Ieperen, W., Joosten, M. H. A. J., and van Kan, J. A. L. 2021. Red light imaging for programmed cell death visualization and quantification in plant–pathogen interactions. Mol. Plant Pathol. 22:361–372.

Li, B., and Fields, S. 1993. Identification of mutations in p53 that affect its binding to SV40 large T antigen by using the yeast two-hybrid system. FASEB J. 7:957–963.

Liebrand, T. W. H., van den Berg, G. C. M., Zhang, Z., Smit, P., Cordewener, J. H. G., America, A. H. P., Sklenar, J., Jones, A. M. E., Tameling, W. I. L., Robatzek, S., Thomma, B. P. H. J., and Joosten, M. H. A. J. 2013. Receptor-like kinase SOBIR1/EVR interacts with receptor-like proteins in plant immunity against fungal infection. Proc. Natl. Acad. Sci. U.S.A. 110:10010–10015.

Liebrand, T. W. H., van den Burg, H. A., and Joosten, M. H. A. J. 2014. Two for all: receptor-associated kinases SOBIR1 and BAK1. Trends Plant Sci. 19:123–132.

Ling, H., Fu, X., Huang, N., Zhong, Z., Liu, T., Cui, H., and Que, Y. 2025. A sugarcane smut fungus effector hijacks plant vacuolar sorting receptor-mediated trafficking to evade host immune detection. Plant, Cell & Environment 48:5271–5289.

Maqbool, A., Saitoh, H., Franceschetti, M., Stevenson, C., Uemura, A., Kanzaki, H., Kamoun, S., Terauchi, R., and Banfield, M. 2015. Structural basis of pathogen recognition by an integrated HMA domain in a plant NLR immune receptor. eLife 4.

Marshall, R. S., and Vierstra, R. D. 2018. Autophagy: the master of bulk and selective recycling. Annu. Rev. Plant Biol. 69:173–208.

Naour-Vernet, M. L., Lahfa, M., Maidment, J. H. R., Padilla, A., Roumestand, C., de Guillen, K., Kroj, T., and Césari, S. 2025. Structure-guided insights into the biology of fungal effectors. New Phytol. 246:1460–1477.

Ortiz, D., de Guillen, K., Cesari, S., Chalvon, V., Gracy, J., Padilla, A., and Kroj, T. 2017. Recognition of the *Magnaporthe oryzae* effector AVR-Pia by the decoy domain of the rice NLR immune receptor RGA5. Plant Cell 29:156–168.

Ose, T., Oikawa, A., Nakamura, Y., Maenaka, K., Higuchi, Y., Satoh, Y., Fujiwara, S., Demura, M., Sone, T., and Kamiya, M. 2015. Solution structure of an avirulence protein, AVR-Pia, from *Magnaporthe oryzae*. J. Biomol. NMR 63:229–235.

Petre, B., Contreras, M. P., Bozkurt, T. O., Schattat, M. H., Sklenar, J., Schornack, S., Abd-El-Haliem, A., Castells-Graells, R., Lozano-Durán, R., Dagdas, Y. F., Menke, F. L. H., Jones, A. M. E., Vossen, J. H., Robatzek, S., Kamoun, S., and Win, J. 2021. Host-interactor screens of *Phytophthora infestans* RXLR proteins reveal vesicle trafficking as a major effector-targeted process. Plant Cell 33:1447–1471.

Pettersen, E. F., Goddard, T. D., Huang, C. C., Meng, E. C., Couch, G. S., Croll, T. I., Morris, J. H., and Ferrin, T. E. 2021. UCSF ChimeraX: structure visualization for researchers, educators, and developers. Protein Sci. 30:70–82.

Postma, J., Liebrand, T. W. H., Bi, G., Evrard, A., Bye, R. R., Mbengue, M., Kuhn, H., Joosten, M. H. A. J., and Robatzek, S. 2016. Avr4 promotes Cf-4 receptor-like protein association with the BAK1/SERK3 receptor-like kinase to initiate receptor endocytosis and plant immunity. New Phytol. 210:627–642.

Rocafort, M., Bowen, J. K., Hassing, B., Cox, M. P., McGreal, B., De La Rosa, S., Plummer, K. M., Bradshaw, R. E., and Mesarich, C. H. 2022. The *Venturia inaequalis* effector repertoire is dominated by expanded families with predicted structural similarity, but unrelated sequence, to avirulence proteins from other plant-pathogenic fungi. BMC Biol. 20:246.

Rocafort, M., Srivastava, V., Bowen, J. K., Díaz-Moreno, S. M., Guo, Y., Bulone, V., Plummer, K. M., Sutherland, P. W., Anderson, M. A., Bradshaw, R. E., and Mesarich, C. H. 2023. Cell wall carbohydrate dynamics during the differentiation of infection structures by the apple scab fungus, *Venturia inaequalis*. Microbiol. Spectr. 11.

Ron, M., and Avni, A. 2004. The Receptor for the Fungal Elicitor Ethylene-Inducing Xylanase Is a Member of a Resistance-Like Gene Family in Tomato. Plant Cell. 16:1604–1615

Sannier, M., Benmamar, S., Collemare, J., Lemaire, C., Caffier, V., Duplaix, C., Shiller, J., Mathieu-Bégné, E., Vayssières, A., Schouten, H. J., Mesarich, C. H., Le Cam, B., and Baudin, M. 2025. Ancient MAX effector variants of a fungal pathogen evade apoplastic immunity in apple. bioRxiv. doi:10.64898/2025.12.25.695362.

Schindelin, J., Arganda-Carreras, I., Frise, E., Kaynig, V., Longair, M., Pietzsch, T., Preibisch, S., Rueden, C., Saalfeld, S., Schmid, B., Tinevez, J.-Y., White, D. J., Hartenstein, V., Eliceiri, K., Tomancak, P., and Cardona, A. 2012. Fiji: an open-source platform for biological-image analysis. Nat. Methods 9:676–682.

Schouten, H. J., Brinkhuis, J., van der Burgh, A., Schaart, J. G., Groenwold, R., Broggini, G. A. L., and Gessler, C. 2014. Cloning and functional characterization of the Rvi15 (Vr2) gene for apple scab resistance. Tree Genet. Genomes 10:251–260.

Sehnal, D., Bittrich, S., Deshpande, M., Svobodová, R., Berka, K., Bazgier, V., Velankar, S., Burley, S. K., Koča, J., and Rose, A. S. 2021. Mol* Viewer: modern web app for 3D visualization and analysis of large biomolecular structures. Nucleic Acids Res. 49–W437.

Steiner, U., and Oerke, E.-C. 2024. The hemibiotrophic apple scab fungus *Venturia inaequalis* induces a biotrophic interface but lacks a necrotrophic stage. J. Fungi 10:831.

Stotz, H. U., Mitrousia, G. K., de Wit, P. J. G. M., and Fitt, B. D. L. 2014. Effector-triggered defence against apoplastic fungal pathogens. Trends Plant Sci. 19:491–500.

Sun, Y., Wang, Y., Zhang, X., Chen, Z., Xia, Y., Wang, L., Sun, Y., Zhang, M., Xiao, Y., Han, Z., Wang, Y., and Chai, J. 2022. Plant receptor-like protein activation by a microbial glycoside hydrolase. Nature 610:335–342.

Švara, A., Sun, H., Fei, Z., and Khan, A. 2024. Chromosome-level phased genome assembly of ‘Antonovka’ identified candidate apple scab-resistance genes highly homologous to *HcrVf2* and *HcrVf1* on linkage group 1. G3 Genes Genomes Genetics 14.

Sweigard, J. A., Carroll, A. M., Kang, S., Farrall, L., Chumley, F. G., and Valent, B. 1995. Identification, cloning, and characterization of PWL2, a gene for host species specificity in the rice blast fungus. Plant Cell 7:1221–1233.

Thomas, C. M., Vos, P., Zabeau, M., Jones, D. A., Norcott, K. A., Chadwick, B. P., and Jones, J. D. G. 1995. Identification of amplified restriction fragment polymorphism (AFLP) markers tightly linked to the tomato Cf-9 gene for resistance to *Cladosporium fulvum*. Plant J. 8:785–794.

van der Hoorn, R. A. L., Laurent, F., Roth, R., and de Wit, P. J. G. M. 2000. Agroinfiltration is a versatile tool that facilitates comparative analyses of Avr9/Cf-9-induced and Avr4/Cf-4-induced necrosis. Mol. Plant-Microbe Interact. 13:439–446.

Vinatzer, B. A., Patocchi, A., Gianfranceschi, L., Tartarini, S., Zhang, H.-B., Gessler, C., and Sansavini, S. 2001. Apple contains receptor-like genes homologous to the *Cladosporium fulvum* resistance gene family of tomato with a cluster of genes cosegregating with Vf apple scab resistance. Mol. Plant-Microbe Interact. 14:508–515.

Weber, E., Engler, C., Gruetzner, R., Werner, S., and Marillonnet, S. 2011. A modular cloning system for standardized assembly of multigene constructs. PLoS ONE 6.

Wu, C.-H., Adachi, H., de la Concepcion, J. C., Castells-Graells, R., Nekrasov, V., and Kamoun, S. 2020. NRC4 gene cluster is not essential for bacterial flagellin-triggered immunity. Plant Physiol. 182:455–459.

Wulff, B. B. H., Thomas, C. M., Smoker, M., Grant, M., and Jones, J. D. G. 2001. Domain swapping and gene shuffling identify sequences required for induction of an Avr-dependent hypersensitive response by the tomato Cf-4 and Cf-9 proteins. Plant Cell 13:255–272.

Zhang, L., Kars, I., Essenstam, B., Liebrand, T. W. H., Wagemakers, L., Elberse, J., Tagkalaki, P., Tjoitang, D., van den Ackerveken, G., and van Kan, J. A. L. 2014. Fungal Endopolygalacturonases Are Recognized as Microbe-Associated Molecular Patterns by the Arabidopsis Receptor-Like Protein RESPONSIVENESS TO BOTRYTIS POLYGALACTURONASES1. Plant Physiol. 164:352–364

Zönnchen, J., Gantner, J., Lapin, D., Barthel, K., Eschen-Lippold, L., Erickson, J. L., Villanueva, S. L., Zantop, S., Kretschmer, C., Joosten, M. H. A. J., Parker, J. E., Guerois, R., and Stuttmann, J. 2022. EDS1 complexes are not required for PRR responses and execute TNL-ETI from the nucleus in *Nicotiana benthamiana*. New Phytol. 236:2249–2264.

