## Supplementary material for "Molecular basis of AvrRvi6 recognition and escape from *Rvi6*-mediated immunity": Figure S

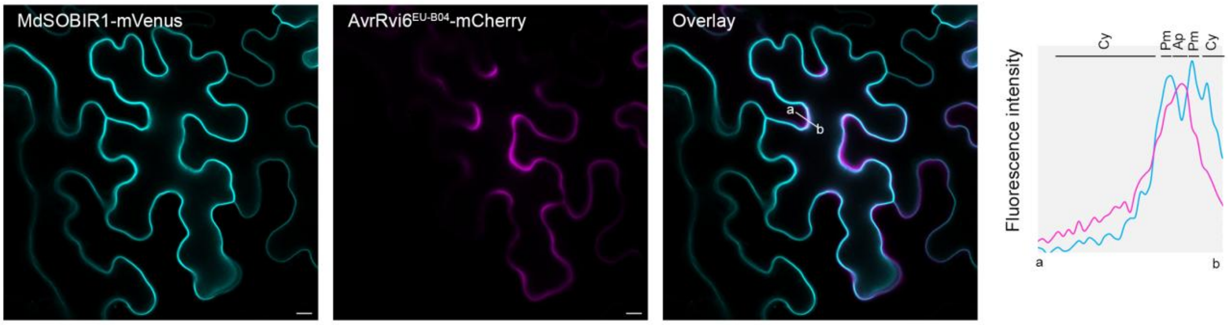


Figure S1: MdSOBIR1 is localized at the plasma membrane in *N. benthamiana.* Subcellular localization of MdSOBIR1, fused to an mVenus fluorescent tag, in *N. benthamiana* leaf epidermal cells at 3 days post-infiltration (dpi) analyzed by laser-scanning confocal microscopy. MdSOBIR1 was co-expressed with the apoplastic avirulent AvrRvi6$^{EU\text{-}B04}$ carrying the plant PR1α signal peptide and a C-terminal mCherry tag (PR1αSP-AvrRvi6ΔSP-mCherry magenta/red), both under a double 35S promoter. MdSOBIR1 localizes to the membrane. The avirulent variant EU-B04 localizes predominantly to the extracellular space (apoplast). Scale bars =10 µm. Signal intensity from point "a" to "b" is shown in a plot. Experiment was repeated at least wice.

```
      19       28        38        48        58        68        78        88
0301  YSCCFEVLGQKDVATGVFANGGVFTWAPRTDCIIEINTNAESCPGWRWRYLSGTSCKTLGLPLAYLGTAPRSQCN
2444  YSCCFEVLGQRDVATGVFANGGVFTWAPRTDCIIEINTNAESCPGWRWRYLSGTSCKTLGLPLAYLGTAPRSQCN
2897  YSCCFEVLGQREVATGVFANGGVFTWAPRTDCIIEINTNAESCPGWRWRYLSGTSCKTLGLPLAYLGTAPRSQCN
1488  YSCCFEVLGQKDVATGVFANGGVFTWAPRTDCIIEINTNAESCPGWRWRYLPGTSCKSLGLPLAYLGTAPRSQCN
2499  YSCCFEVLGQKDVATGVFANGGVFTWAPRTDCIIEINTNAESCSGWRWRYLSGTSCKTLGLPLAYLGTAPRSQCN
2917  YSSCFEVLGQKDVATGVFANGGVFTWAPRTDCIIEINTNAESCPGWRWRYLSGTSCKTLGLPLAYLGTAPRSQCN
2407  YSCCFEVLGQKDVATGVFANGGVFTWAPRTDCIIEINANAESCPGWRWRYLSGTSCKTLGLPLAYLGTAPRSQCN
2484  YSCCFEVLGQRDVATGVFANGGVFTWAPRTDCIIEINTNAESCPGWRWRYLSGTSCKSLGLPLAYLGTAPRSQCN
2230  YSCCFEVLGQKDVATGVFANGGVFTWAPRTDCIIEINANAESCPGWRWRYLPGTSCKSLGLPLAYLGTAPRSQCN
1395  YSCCFEVLGQKHVATGVFANGGVFTWAPRTDCIIEINANAESCPGWRWRYLSGTSCKTLGLPLAYLGTAPRSQCN
2484  YSCCFEVLGQKDVATGVFANGGVFTWAPRTDCIIEINANAESCPGWRWRYLSGTSCKSLGLPLAYLGTAPRSQCN
2482  YSCCFEVLGQKDVATGVFANGGVFTWAPRTDCIIEINANAESCSGWRWRYLSGTSCKSLGLPLAYLGTAPRSQCN
EUB04 YSCCFEVLGQRDVATGVFANGGVFTWAPRTDCIIEINTNAESCSGWRWRYLSGTSCKSLGLPLAYLGTAPRSQCN
2480  YSCCFEVLGQRDVATGVFANGGVFTWAPRTDCIIEINANAESCSGWRWRYLSGTSCKSLGLPLAYLGTAPRSQCN
1402  YSCCFEVLGQKDVATGVFANGGVFTWAPSTDCIIEINANAESCSGWRWRYLSGTSCKTLGLPLAYLGTAPRSQCN
1153  YSCCFEVLGQKDVATGVFANGGVFTWAPRTDRIIEINTNAESCSGWRWRYLSGTSCKTLGLPLAYLGTAPRSQCN
1459  YSCCFEVLGQRDVATGVFANGGVFTWAPRTDCIIEINANAESCPGWRWRYLSGTSCKSLGLPLAYLGTAPRSQCN
1180  YSCCFEVLGQKDVATGVSANGGVFTWAPRTDCIIEINTNAESCSGWRWRYLSGTSCKTLGLPLAYLGTAPRSQCN
2466  YSCCFEVLGQKDVATGVFANGGVFTWAPLTDCIIEINANAESCSGWRWRYLSGTSCKTLGLPLAYLGTAPRSQCN
```

Figure S2: AvrRvi6 alleles sequence alignment. All AvrRvi6 alleles studied, except for the truncated AvrRvi6$^{2828}$ allele have been aligned. polymorphisms present in both virulent and avirulent alleles are highlighted in yellow, those present only in virulent alleles are shown in red. Virulent allele names are written in red, avirulent ones in black.

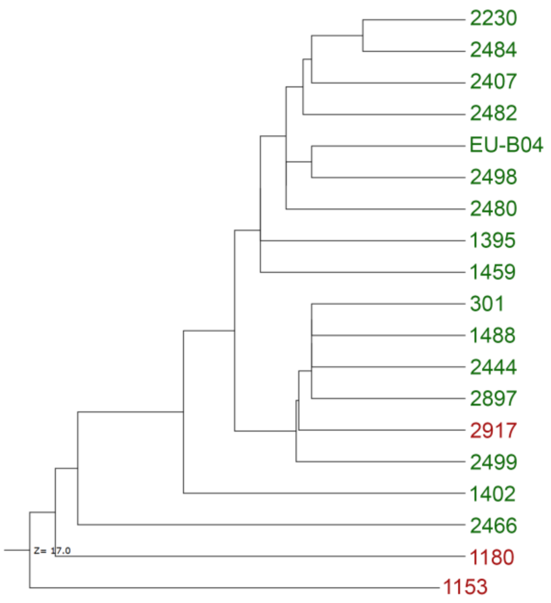


<u>Figure S3:</u> <u>Structural dendrogram of AvrRvi6 alleles</u>. All AvrRvi6 alleles studied, except for the truncated AvrRvi6$^{2828}$ allele have been 3D modeled and structurally aligned. Avirulent allele names are written in green and virulent ones in red.

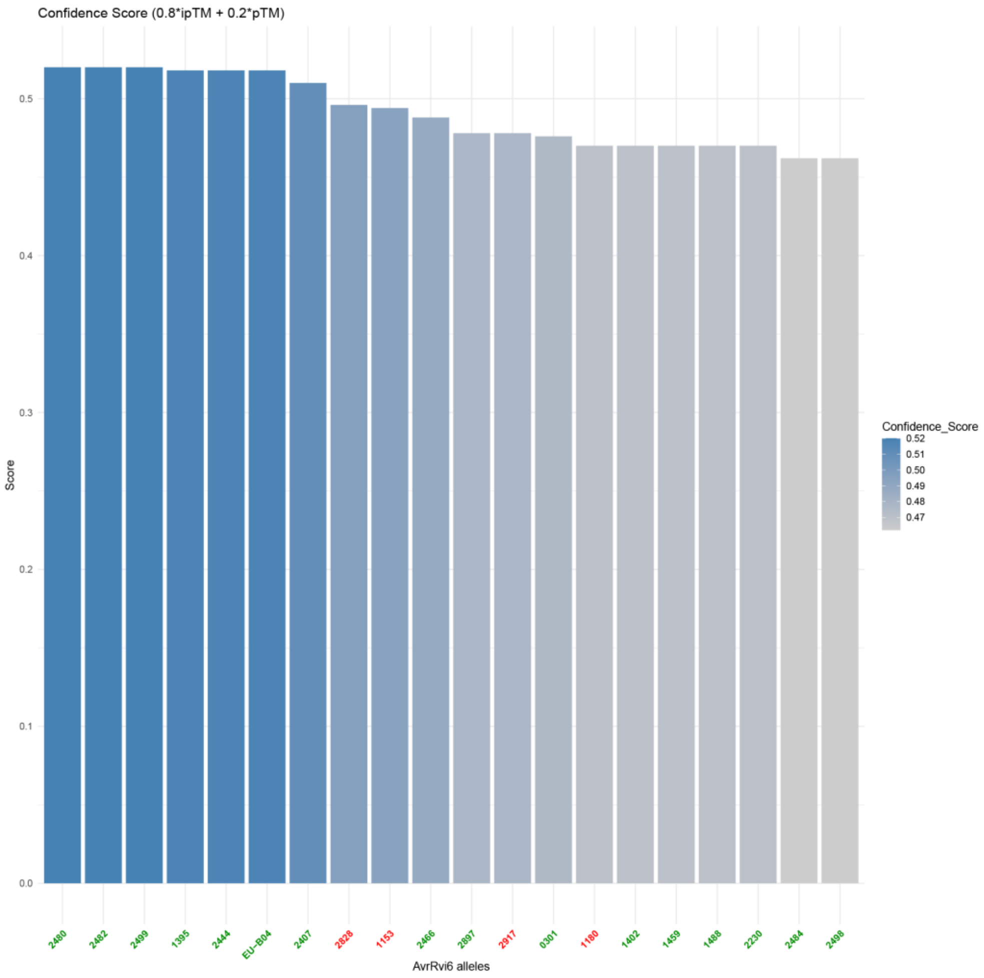


Figure S4: Confidence score analysis of Rvi6-AvrRvi6 complexes. Complex model confidence was determined using the weighted interface metric: 0.8 ipTM + 0.2pTM (Evans et al. 2021). Virulent allele names are written in red, avirulent ones in green.

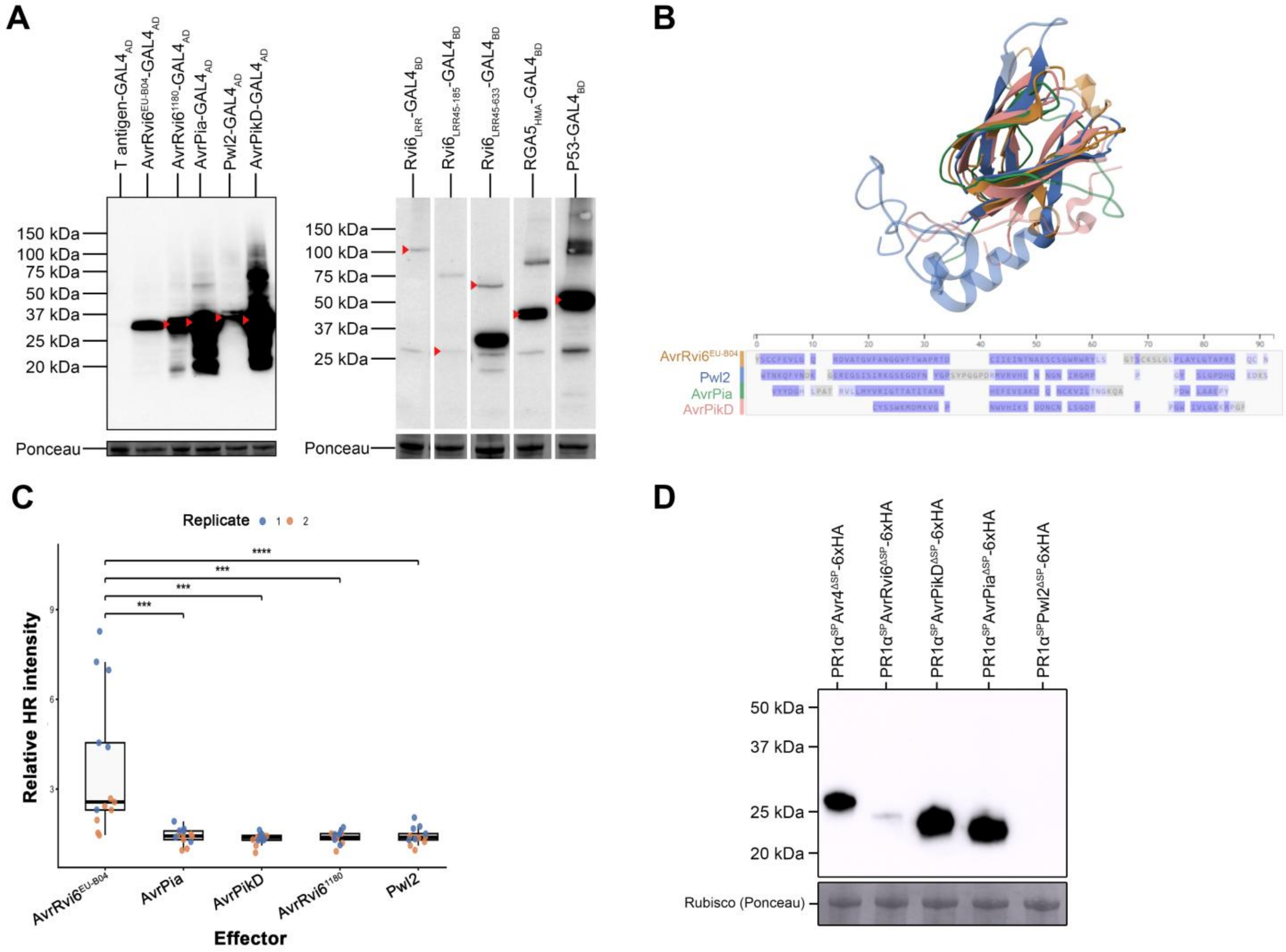


Figure S5: Rvi6$_{LRR45–633}$ associates with AvrRvi6 and some other MAX effectors in yeast but only AvrRvi6 is recognized triggering an HR in *N. benthamiana.* A) Immunoblot analysis confirming the steady-state expression of proteins involved in the Y2H experiment (Figure 6). B) Structural alignment of MAX effectors tested in Y2H experiment with the reference AvrRvi6$^{EU\text{-}B04}$. C) Functional assessment of HR induction in *N. benthamiana* leaves upon transient co-expression of Rvi6 with different MAX effectors. D) Immunoblot analysis confirming the steady-state expression of effectors expressed in *N. benthamiana* in (D).

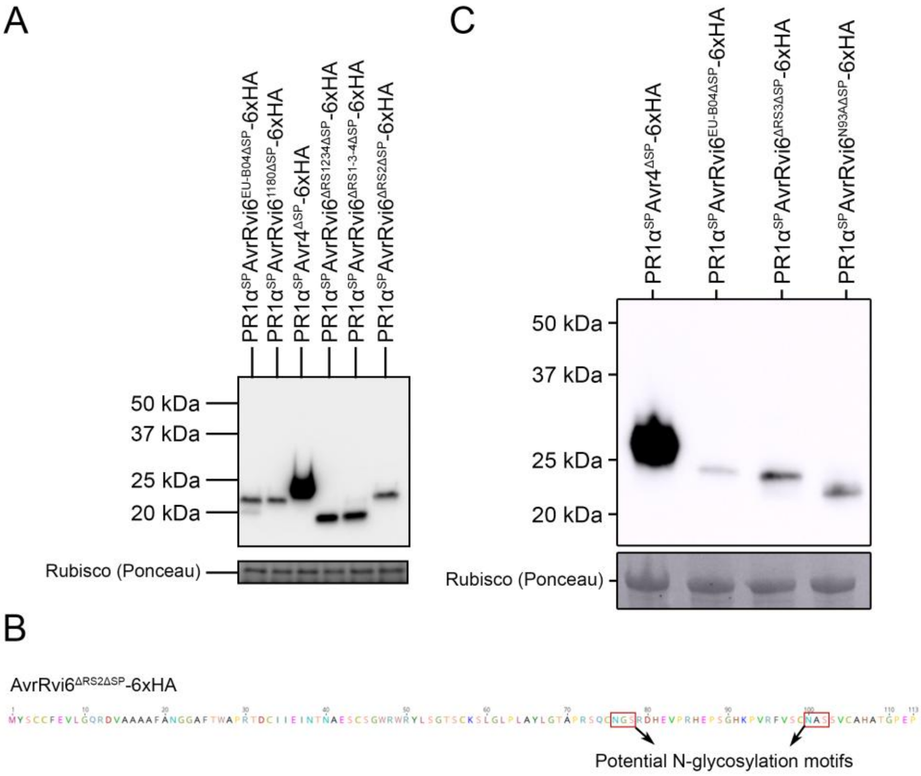


<u>Figure S6:</u> <u>AvrRvi6 recognition-site-mutants are well expressed in *N. benthamiana*.</u> A) Immunoblot analysis confirming the steady-state expression of AvrRvi6$^{EU\text{-}B04}$, AvrRvi6$^{1180}$ and the AvrRvi6 mutants (AvrRvi6$^{\Delta RS1234}$, AvrRvi6$^{\Delta RS134}$ and AvrRvi6$^{\Delta RS2}$), in *N. benthamiana* during HR induction tests (Figure 7C). B) Protein sequence of AvrRvi6$^{\Delta RS2\Delta SP}$-6xHA with indicated potential N-glycosylation motifs. C) Immunoblot analysis confirming the steady-state expression of AvrRvi6$^{\Delta RS3}$, AvrRvi6$^{N93A}$ , AvrRvi6$^{EU\text{-}B04}$ in *N. benthamiana* during HR induction tests (Figure 7D) Avr4 is used as a positive control.
